# Pervasive relaxed selection in termite genomes

**DOI:** 10.1101/2023.11.01.565207

**Authors:** Kyle M. Ewart, Simon Y. W. Ho, Al-Aabid Chowdhury, Frederick R. Jaya, Yukihiro Kinjo, Juno Bennett, Thomas Bourguignon, Harley A. Rose, Nathan Lo

## Abstract

The genetic changes that enabled the evolution of eusociality have long captivated biologists. In recent years, attention has focussed on the consequences of eusociality on genome evolution. Studies have reported higher molecular evolutionary rates in eusocial hymenopteran insects compared with their solitary relatives. To investigate the genomic consequences of eusociality in termites, we sequenced genomes from three of their non-eusocial cockroach relatives. Using a phylogenomic approach, we found that termite genomes experienced lower rates of synonymous mutations than those of cockroaches, possibly as a result of longer generation times. We identified higher rates of nonsynonymous mutations in termite genomes than in cockroach genomes, and identified pervasive relaxed selection in the former (24–31% of the genes analysed) compared with the latter (2–4%). We infer that this is due to a reduction in effective population size, rather than gene-specific effects (e.g., indirect selection of caste-biased genes). We found no obvious signature of increased genetic load in termites, and postulate efficient purging at the colony level. Additionally, we identified genomic adaptations that may underpin caste formation, such as genes involved in post-translational modifications. Our results provide insights into the evolution of termites and the genomic consequences of eusociality more broadly.

## 1. Introduction

Social insects represent only ∼2% of all insect species but have an enormous ecological impact, representing over half of all insect biomass [1]. Eusocial insect colonies comprise highly specialized castes, whereby some members of the colony reproduce while the workers and soldiers focus on tasks such as foraging, defence, brood care, and nest construction [2]. These considerably different behaviours arise from the same genome. The behavioural and genetic changes that enabled the evolution of eusociality, and that have facilitated the ecological dominance of these insects, have long interested biologists.

Most studies on the evolution of eusociality have focussed on the genomic features that underpin this trait. However, in recent years, the consequences of eusociality on genome evolution have also become the subject of active research [3–6]. A key parameter of interest is the molecular evolutionary rates of eusocial organisms compared with their solitary relatives [7,8]. Two main hypotheses have been put forward in relation to the consequences of eusociality on evolutionary rates. The first hypothesis posits that the presence of very few reproductive individuals within colonies should result in eusocial insects having small effective population sizes (*N_e_*). This would be expected to increase the strength of genetic drift in eusocial lineages, leading to higher evolutionary rates [3,5,9–12] (but see [13]). The second hypothesis states that the specific expression of genes in different castes should reduce pleiotropy and distinct selection pressures acting on these genes (i.e., ‘indirect selection’), leading to the reduced efficacy of selection [6,7,14,15]. Moreover, genes under relaxed selection in eusocial ancestors might more readily adopt caste-biased expression, hence genes that are caste-specific in eusocial species might have elevated evolutionary rates in both eusocial and non-eusocial taxa [8].

Studies of hymenopteran social insects have identified elevated rates of genome-wide evolution [5], as well as elevated rates of evolution in differentially expressed caste-specific genes [7,8], supporting both of the aforementioned hypotheses. However, the relative contributions of reduced *N_e_*and differential gene expression to these elevated rates remain unclear. Further, there has been limited use of phylogenomic approaches to investigate rates of genomic evolution in eusocial insects and their relatives, despite the availability of methods for examining differing molecular evolutionary forces across large numbers of loci. Phylogenomic methods can help to elucidate the timing of shifts in genomic rates, characterize signatures of selection associated with rate changes, and identify the specific genes affected. Further, most previous studies have focussed on hymenopterans, with relatively little attention given to genome evolution in other eusocial insects.

An important eusocial insect group with significant ecological impacts is the termites. Their evolution of eusociality from cockroach ancestors [16–20], and their diploid chromosome makeup (in contrast to the haplodiploid eusocial species in Hymenoptera), makes them a valuable comparative model system. However, past examinations of genome-wide molecular evolution in termites have typically been based on a single termite representative [3,5], or have lacked comparisons with cockroach relatives [6]. Although a number of termite genomes have been sequenced [21–24], genomes are available for only two cockroach species, the pests *Periplaneta americana* (Blattidae [25]) and *Blattella germanica* (Ectobiidae [22]). A lack of genomic resources hinders our ability to perform in-depth comparisons of termites and cockroaches to examine the consequences of eusociality on genome evolution.

Here, we facilitate comparative phylogenomic analyses of Blattodea by sequencing the genomes of three Australian species from Blaberidae, the most speciose cockroach family. *Panesthia cribrata* (subfamily Panesthiinae) nests in and feeds on rotting wood, but displays minimal social behaviour. *Geoscapheus dilatatus* and *Neogeoscapheus hanni* (subfamily Geoscapheinae) feed on dried leaf litter and form burrows in the soil up to a metre deep, in which offspring remain with their mothers for several months. The inclusion of *P. cribrata* in our analyses allowed us to control for the effects of wood nesting and wood feeding on genome evolution, in the absence of eusocial behaviour.

We utilize these new data in conjunction with previously generated genomes from Blottodea to achieve three main objectives: (1) increase our understanding of patterns of selection across termite and cockroach genomes; (2) to evaluate factors that have been posited to influence molecular evolutionary rates in eusocial organisms, such as a reduced *N*_e_ and indirect selection of caste-biased genes; and (3) to identify genomic adaptations that underpin the evolution of eusociality in termites. We expect that this study will shed light on the evolution of termites and eusociality more broadly.

## 2. Results and discussion

### (a) Genome assemblies and orthologue assignment

We produced novel *de novo* genome assemblies for *G. dilatatus*, *N. hanni*, and *P. cribrata* using a combination of high-throughput sequencing approaches (Table S1; Figure S1). The genome quality for *G. dilatatus* was comparable to that of the well-studied pest cockroach species *Periplaneta americana* and *Blattella germanica* (Table S2 [22,25]), while genome completeness and continuity for *P. cribrata* and *N. hanni* were lower (Table 1). Annotated protein-coding genes were BLASTed against the insect protein database, resulting in an average sequence identity of >65% for each species (see Figure S2 for annotation QC results). Most annotated genes (91.7%) were assigned to orthogroups, and relatively few were found to be species-specific (Figure S3). Of the 24,455 orthogroups, 1491 were single-copy and present in all species; we used these single-copy orthologues in the subsequent evolutionary analyses.

**Table 1.** Genome assembly and annotation metrics. *Ab initio* predictions that had no BLAST hits were not included in the numbers of annotated genes.

| Species | Genome length | N50 | No. of contigs | Complete BUSCOs | Fragmented BUSCOs | Missing BUSCOs | Duplicated BUSCOs | No. of annotated genes |
| --- | --- | --- | --- | --- | --- | --- | --- | --- |
| *G. dilatatus* | 1.70 Gb | 3.2 Mb | 84,284 | 93.9% | 4.8% | 1.3% | 0.7% | 30,006 |
| *N. hanni* | 1.68 Gb | 11.4 kb | 270,019 | 69.0% | 19.5% | 11.5% | 0.7% | 38,314 |
| *P. cribrata* | 1.78 Gb | 298 kb | 204,297 | 85.0% | 10.9% | 4.1% | 1.8% | 39,852 |

### (b) Pervasive relaxed selection in termite genomes

By investigating patterns of *d_N_* and *d_S_*, we found evidence of pervasive relaxed selection on the termite terminal branches of the phylogeny. Our analyses included estimates of *d_N_* and *d_S_* for all single-copy orthologues in nine examined taxa: the three newly sequenced cockroach species (*Geoscapheus dilatatus*, *Neogeoscapheus hanni* and *Panesthia cribrata*), two pest cockroach species (*Periplaneta americana* and *Blattella germanica*), three termite species (*Cryptotermes secundus*, *Coptotermes formosanus* and *Zootermopsis nevadensis*), and one outgroup species (*Laupala kohalensis*).

We applied the free-ratio model in CODEML [26], allowing a separate *d_N_* and *d_S_* on each branch, and found that median *d_N_* rates were found to be elevated on the termite branches (excluding the termite stem; median = 4.19×10^-4^) compared with the cockroach branches (excluding the Blattodea stem; median = 2.85×10^-4^). Conversely, the *d_S_* rate was found to be lower on the termite branches (median = 2.66×10^-3^) than on the cockroach branches (median = 3.46×10^-3^). The median *d_N_*/*d_S_*for the termite branches was 0.149, while that for cockroach branches was 0.081. Separate analyses of *d_N_*/*d_S_*utilizing a three-ratio model (allowing a separate ratio for the termite clade) was preferred over a two-ratio model (allowing one ratio for the outgroup and one for the ingroup) for 75.7% of the analysed orthologues (*p* < 0.05; 45.1% after applying a Holm-Bonferroni correction). The distribution of *d_N_*/*d_S_*on termite branches in this analysis was wider and had a much higher mean than that for the cockroach branches (Figure 1a). In 90.8% of the analysed orthologues, the termite *d_N_*/*d_S_* was higher than the cockroach *d_N_*/*d_S_* (significantly higher for 73% when applying *p* < 0.05, and 44.5% after applying a Holm-Bonferroni correction).

**Figure 1.**
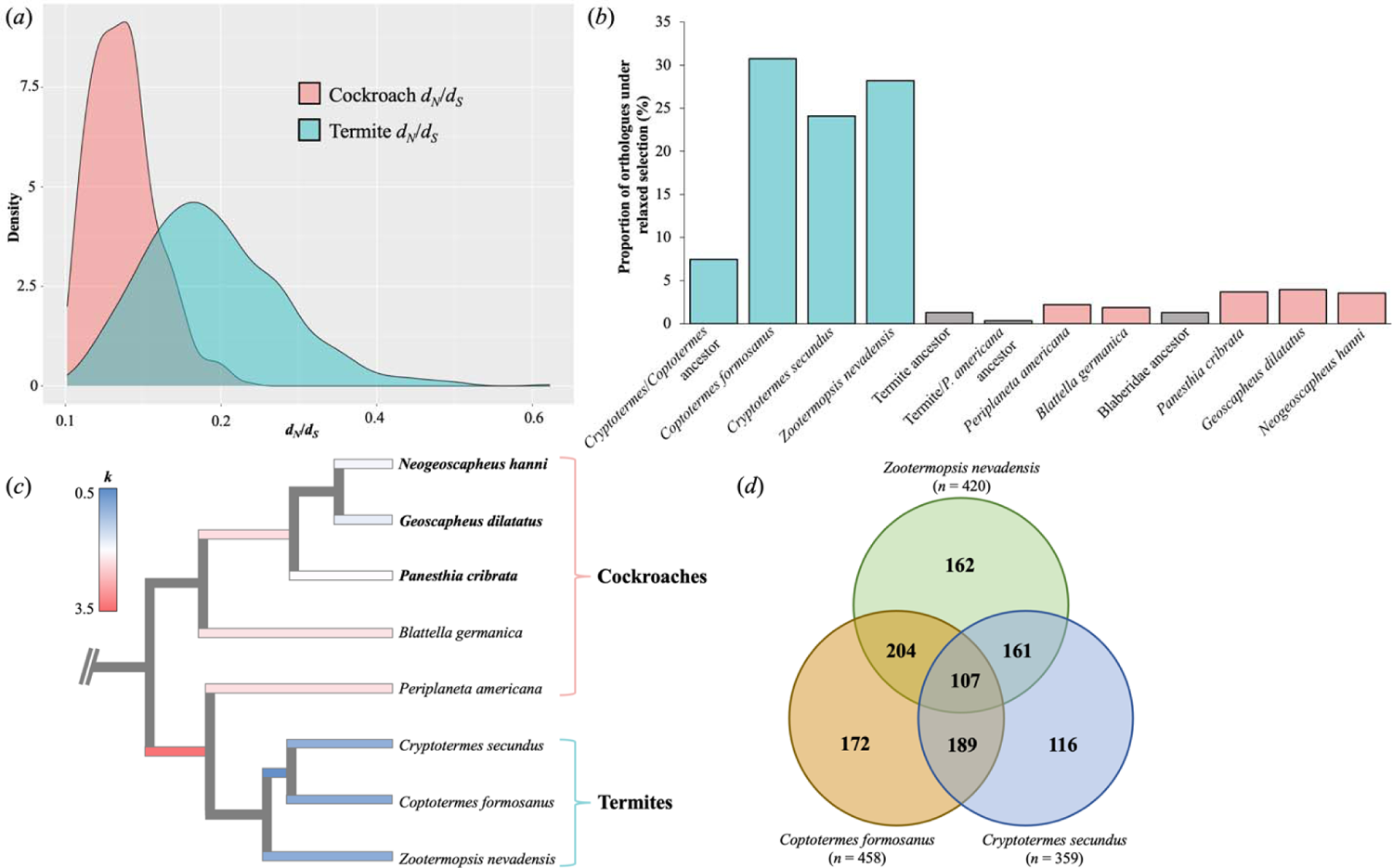
(a) Density plot for the 3-ratio *d_N_*/*d_S_* model for the cockroaches and termites. (b) Proportion of genes under significant relaxed selection for each of the focal branches based on the RELAX analyses (after Holm-Bonferroni correction). (c) Phylogenetic tree of analysed termite and cockroach genomes (genomes generated in this study in bold), with focal branches in the RELAX analyses coloured by their median relaxed selection parameter (*k*), whereby *k* < 1 indicates relaxed selection and *k* > 1 indicate intensified selection (the outgroup has been pruned from the tree). (d) Venn diagram displaying overlapping orthologues under relaxed selection on the termite terminal branches.

To investigate further the differences in *d_N_*, *d_S_*, and *d_N_*/*d_S_* between termite and cockroach branches, we tested for relaxed selection using the ‘RELAX’ method [27], implemented in HyPhy [28,29]. We detected a large proportion of the single-copy orthologues under relaxed selection on the termite terminal branches (24.09–30.74%) compared with the cockroach terminal branches (1.88–3.96%) (after a Holm-Bonferroni correction for testing multiple branches; Figure 1b). When we analysed an audited subset of single-copy orthologues (i.e., gene alignments that were manually inspected and edited if required), we obtained similar proportions (Table S3). In relaxed selection analyses, a significant result of *k* < 1 is indicative of a reduction in selection strength, while *k* > 1 indicates an intensification of selection strength. Median *k* values were found to be the lowest on termite branches, and well below one (Figure 1c).

### (c) Relaxed selection in termite genomes is better explained by reduced *N_e_* rather than indirect selection

We evaluated two main scenarios that may explain the pervasive selection in termite genomes: 1) a reduction in *N*_e_, and 2) indirect selection. A reduction in *N_e_* would have a genome-wide effect, whereby the strength of genetic drift increases for every gene. Conversely, indirect selection would have gene-specific effects. Only caste-specific genes are expected to experience reduced selective constraints, as they are only expressed in a subset of the colony (e.g., worker-biased genes are predominantly expressed by workers, not queens). We note that a niche- or functional-specialization scenario would also confer gene-specific effects (elaborated in section ‘d’ below).

To investigate whether indirect selection underpinned signatures of relaxed selection in our data set, we identified and analysed 311 genes in our single-copy orthologue data set that had previously been identified to display either unbiased or caste-biased differential expression [6,30] (noting that caste-biased genes in *Coptotermes formosanus* were not identified in these studies). We found no significant difference between the proportion of caste-biased (33.3% for *Zootermopsis nevadensis* and 24.4% for *Cryptotermes secundus*) and unbiased (25.3% for *Zootermopsis nevadensis* and 20% for *Cryptotermes secundus*) orthogroups under relaxed selection when compared with proportions in the full single-copy orthologue data set (chi-squared tests, *p* = 0.29 for *Zootermopsis nevadensis* and *p* = 1 for *Cryptotermes secundus* caste-biased genes; Table S4). Similar patterns were found for genes only expressed in one caste (Table S4). Further, if indirect selection were the main driver of these patterns of relaxed selection, we would expect to see considerably fewer genes affected. Previous RNAseq analyses showed that many caste-biased genes are species specific [6] and/or part of multi-gene families [30]. Such genes were not considered in our single-copy orthologue data set and require further investigation.

The pervasive scale of relaxed selection across the termite genomes is explained better by a reduction in *N_e_* than by gene-specific effects resulting from indirect selection. A reduction in *N_e_*increases the intensity of genetic drift [31], and consequently reduces the efficacy of both negative and positive selection [27,32]. Termites are expected to have a very low *N_e_*relative to their census population size due to the complex social structure of their colonies, the small number of reproductive individuals (typically one king and one queen per colony), and their history of inbreeding within colonies [11,33,34]. Indeed, Romiguier et al. [5] found that the termite *Reticulitermes grassei* had a large reduction in *N_e_*, even compared with other eusocial species, and postulated that this was due to the strict monogamy seen in this taxon.

Although a reduced *N_e_* will theoretically not directly affect the substitution rate at neutral sites [35], a long-term low *N_e_* has been proposed to lead to the evolution of a higher mutation rate [36]. However, we found that *d_S_* values were lower in termites than in cockroaches. This result can be explained by the considerably longer generations of termites (5–10 years) compared with cockroaches (typically <1 year). Although a longer generation time is expected to increase the mutation rate per generation (due to increased cell divisions of gametes, and greater time for DNA damage to accumulate), the per-year mutation rate is expected to decrease [37,38]. However, a reduction of *N_e_* appears to have counteracted this generation-time effect for non-neutral sites in termites (based on observed *d_N_*rates).

### (d) Alternative explanations for relaxed selection in termite genomes

Although reduced *N_e_* is the most likely explanation for the pervasive relaxed selection observed in our study, we are unable to rule out the possibility that indirect selection or other factors contributed to relaxed selection in at least some of the examined orthologues. Although many genes were only under significant relaxed selection on one of the three termite branches (Figure 1d), there were more genes under relaxed selection that overlap in all three termite species (*n* = 107) than expected by chance. Hence, some ‘core genes’ might be under relaxed selection due to other non-random processes, such as indirect selection and/or reduced functional importance in termites. There was no evidence that caste-biased genes were undergoing relaxed selection or accelerated evolution prior to becoming caste-biased, as suggested by previous authors [8,39,40], based on evolutionary rates of the cockroach orthologues (Figures S4, S5) and the minimal relaxed selection on the termite ancestral branch (Figure 1b–c).

The intensity of selection is proportional to the product of *N_e_*and the selection coefficient *s* [35]. Therefore, it is possible to observe pervasive relaxed selection without a reduction in *N_e_*, given a change in *s* that affects many loci. Previous authors have hypothesized that the cooperative interactions of social species can buffer or compensate for the expression of deleterious mutations, particularly in relation to immune responses [41–46]. The creation of a benign environment through social cooperation (e.g., termite mounds) and/or the compensating for the expression of deleterious alleles (e.g., through food allocation) will lead to a reduction in selection pressures for the genes involved [46]. For example, termite social hygiene behaviours, such as allogrooming, might result in relaxation of selection in certain innate components of their immune system [45].

Although there might have been large-scale shifts in *s* across termite genomes since divergence from their subsocial cockroach ancestors, we have several reasons to believe that this is not the primary driver of the patterns of relaxed selection observed. First, said patterns do not appear to be associated with specific habitats. The different termite species analysed occupy different niches: *Coptotermes formosanus* constructs nests and forages in the soil; *Zootermopsis nevadensis* lives and feeds in dead dampwood; and *Cryptotermes secundus* lives and feeds in dead drywood. Likewise, the five sampled cockroach species live in varying habitats, from urban environments to wood and soil burrows. Hence, if pervasive relaxed selection were caused by a reduced *s* due to differing selective pressures across different niches (and not a reduction in *N_e_*), we might expect relaxed selection to be restricted to one or two of the termite species, or observe comparable levels of relaxed selection in some cockroach species (e.g., *Periplaneta americana*, a recently urbanized cockroach species, sister lineage to the termite species in our analysis). Second, if a reduced *s* across many genes were due to the social benefits of group living (e.g., allogrooming), we might expect a larger overlap of orthologues to be under relaxed selection (Figure 1d; but see [43]), and a larger proportion of orthologues under relaxed selection in the termite ancestral branches. Third, given the relatively conserved data set that we analysed (i.e., single-copy orthologues across taxa ∼350 million years divergent [47]), we expect most genes to be functionally important in termites (i.e., a relatively stable *s* across the analysed phylogeny). Hence, a reduction in *s* across >24% of the relatively conserved orthologue data set is unlikely. Fourth, if some of the analysed orthologues were indeed associated with changes in niche or function, we might also expect to see signatures of positive selection in termites; however, this was not the case (Figure 2a). Therefore, although we cannot rule out a pervasive reduction of *s* across the termite genome due to increased social cooperation or niche specialization, or a combination reduced *s* and indirect selection, our results suggest that the observed patterns of relaxed selection are dominated by the genome-wide effects of a reduced *N_e_*, rather than gene-specific effects.

**Figure 2.**
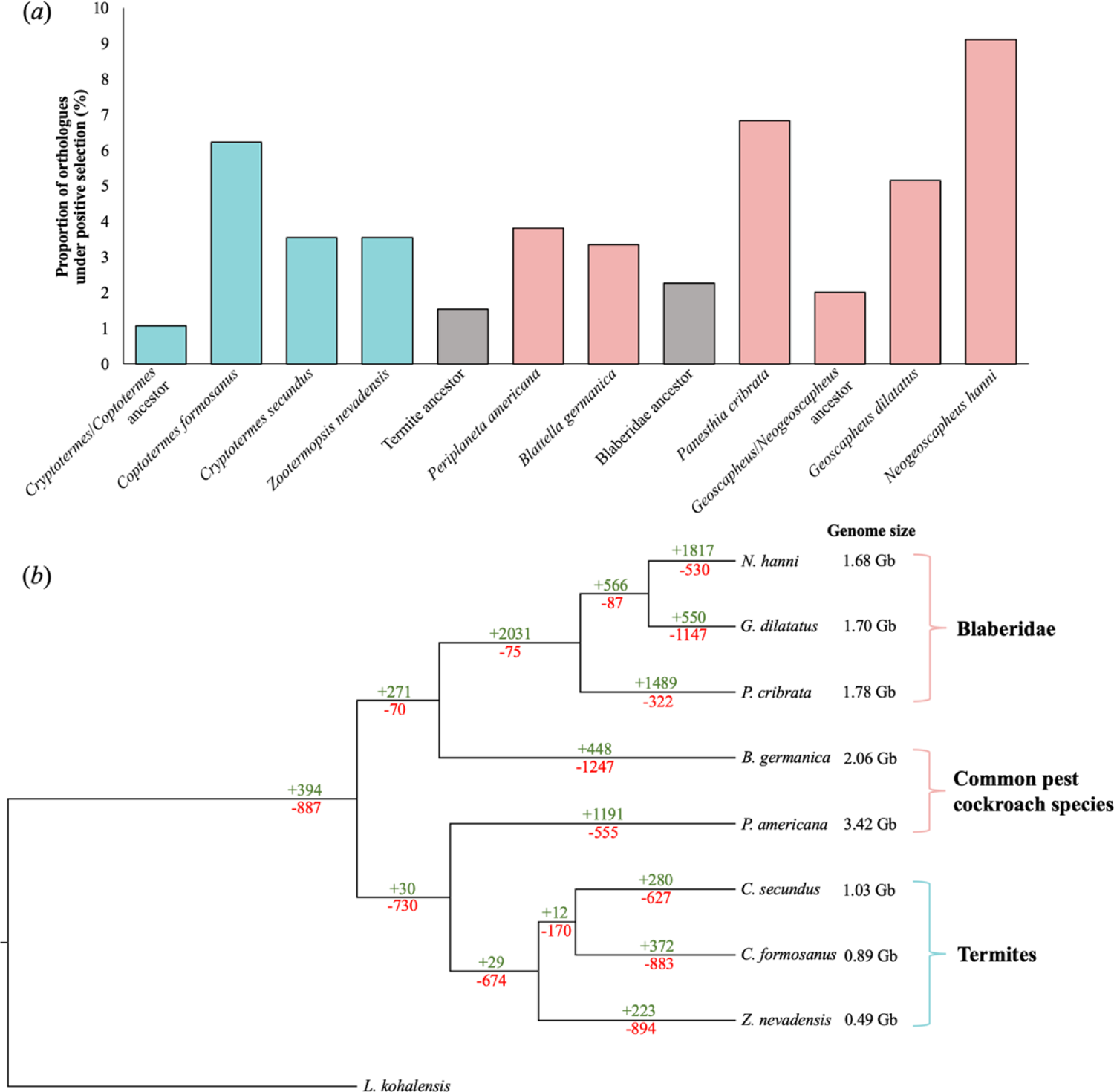
(a) Proportion of orthologues (*n* = 1491) on various branches undergoing positive selection, determined using an aBSREL analyses. (b) Number of gene expansions (green) and contractions (red) on each branch of the cockroach/termite tree. The number of gene expansions/contractions combined results from the small and large orthogroups analysed.

### (e) No obvious signature of increased genetic load in termites

A long-term small *N_e_* and pervasive relaxed selection could result in an accumulation and fixation of deleterious alleles and a high genetic load [3,32,48,49]. Unexpectedly, however, termites do not appear to have a higher genetic load than cockroaches based on the number of derived putative deleterious substitutions (identified using PROVEAN [50]) in 581 insect BUSCOs (i.e., highly conserved genes). Here, we are referring only to ‘realized load’ and not ‘masked load’ [49,51], given that we are comparing amino acid sequences across an order of insects, and not considering heterozygosity. Termites had a higher number of total substitutions per amino acid than the cockroach species (significantly higher for 12/15 pairwise comparisons, and non-significantly higher for 3/15 pairwise comparisons; Figure S6; Table S5). However, the number of inferred deleterious substitutions per amino acid was significantly lower or similar in termites compared with the cockroach species.

Therefore, despite the patterns of pervasive relaxed selection that we identified, there does not appear to be an obvious signature of increased genetic load in termites compared with cockroaches. How then have termites, an ancient and evolutionarily successful group, succeeded in diversifying and adapting to changing environments? An increased genetic load could have been alleviated through purging, which involves the removal of recessive deleterious alleles exposed by inbreeding. Purging could be particularly effective in termites due to a long-term history of inbreeding and variation in heterozygosity generated by inbreeding [52,53]. Moreover, termites could have experienced purging at the colony level. The survival of dispersing reproductive individuals that attempt to form a new colony relies on the survival of their offspring; colony survival hinges on the worker offspring gathering resources, building the nest structure, and raising offspring. Therefore, if considering the colony as the basic unit of the population rather than individual termites (as in [33,54–56], when primary (alate-derived) reproductives disperse and breed, purging could occur by the failure of colonies established by unfit offspring. Indeed, there is evidence that closely related primary reproductives establish colonies less frequently than non-related primary reproductives, as a result of inbreeding depression [57] (but see [58–61]). This hypothesis requires further investigation.

The lack of an obvious increased genetic load in termites may also be due to the insensitivity of the methods we applied, particularly given the complexities of delineating benign and deleterious substitutions. Further investigation into the functional effects of mutations in coding regions (e.g., experimental evidence) may help to characterize the deleteriousness of derived mutations and clarify any differences in genetic load in these species [51]. Further, we acknowledge issues of phylogenetic non-independence in our data set, though we believe that this has a negligible effect on these inferences (detailed in the electronic supplementary material).

### (e) Positive selection and gene expansions/reductions in termite and cockroach genomes

In our analyses of positive selection that allowed a separate *d_N_*/*d_S_*on terminal branches leading to cockroach and termite taxa, we did not detect substantial positive selection in termite genomes (detailed further in the electronic supplementary material). The number of orthologues under positive selection varied between 1.1% and 9.1% across the terminal branches (Figure 2a), with the clade comprising *P. cribrata*, *G. dilatatus*, and *N. hanni* displaying the highest number. Further, no significant differences were found when comparing the proportions of genes under positive selection that had previously been demonstrated to be unbiased or differentially expressed between castes (Table S4).

Previous studies on eusocial species have highlighted the importance of epigenetic mechanisms, particularly DNA methylation, for controlling caste-specific gene expression [22,56,62–67]. Of the 8538 orthogroups that we identified to be displaying gene family size changes (Figure 2b; detailed further in the electronic supplementary material), numerous expanding gene families in the termite lineages were related to protein ubiquitination (Table S7; e.g., E3 ubiquitin-protein ligase, and Ubiquitin carboxyl-terminal hydrolase 45), an important post-translational mechanism, as well as other gene families involved in gene expression regulation (e.g., Histone H4 transcription factor and KAT8 regulatory NSL complex subunit 3). Protein ubiquitination results in protein degradation or modification. Histones are often ubiquitinated, which subsequently alters chromatin structure and gene function, and may therefore influence patterns of DNA methylation [68,69]. Previous studies have demonstrated that these post-translational modifications and histone structure changes influence the development of castes in the honey bee [70–72] and carpenter ant [73,74]. Evidently, various gene expression mechanisms beyond DNA methylation, such as ubiquitination, could be fundamental to the development of castes, and therefore warrant further research.

## 3. Concluding remarks

Incorporating additional genome resources has proven valuable in our comparative genomic analyses of termites and cockroaches. Sequencing and analysing genomes from the sister group of termites, representatives of the wood-feeding genus *Cryptocercu*s, will help to elucidate the ancestral signals of the relaxed selection that we have identified, and provide insights into the evolutionary origins of caste-biased genes. Furthermore, expanding the analyses to include genomes of other eusocial groups (e.g., Hymenoptera) could shed light on convergent patterns of eusocial evolution (e.g., [75]).

Our findings corroborate several previous studies that suggested an association between eusociality and a reduced *N_e_*, resulting in relaxed selection [3,5,12,37]. Although accurate estimation of *N_e_* is particularly challenging for eusocial species [76], integrating such estimates, or relevant proxies (e.g., [5]), could help to clarify the patterns of relaxed selection that we have inferred, and the association between *N_e_* and the evolution of eusociality. This, in turn, would help to illuminate the extent to which shifts in *N_e_* and the emergence of eusociality have shaped genomic evolution in termites.

## 4. Material and methods

### (a) Sample collection and DNA extraction

Samples of *P. cribrata* were collected from North Manly, NSW, in August 2020 by N. Lo. Samples of *G. dilatatus* were collected from Gilgandra, NSW, in September 2019 by H. A. Rose. Samples of *N. hanni* were collected from Mt Molloy, QLD, in January 2020 by J. A. Walker and N. Lo. Muscle tissue was removed from the leg of one adult female of each species for DNA extraction. DNA was extracted using a phenol/chloroform extraction protocol (detailed in the electronic supplementary material).

### (b) DNA sequencing approaches

Three different DNA sequencing approaches were employed to generate the cockroach genomes: linked-read sequencing, chromosomal conformation capture sequencing, and long-read sequencing (Table S1). We used two technologies for linked-read sequencing: Transposase Enzyme-Linked Long-read Sequencing (TELL-Seq™ [77]) and single-tube long fragment read (stLFR [78]). These technologies are based on co-barcoding short-read sequences that derive from the same DNA fragment, providing long-range information to enable assembly of long DNA regions.

We utilized Hi-C sequencing [79], a chromosomal conformation capture sequencing method based on proximity ligation. Hi-C sequencing supports scaffolding of genomic regions that are in close spatial proximity. Hi-C library preparation was performed using the Arima Hi-C Plus kit (Arima, USA) following the manufacturer’s protocol for large animal tissue, using the *DpnII* and *HinfI* enzymes. Preliminary sequencing and subsequent analysis using qc3C [80] was performed to estimate the quantity of Hi-C sequencing required.

We utilized the Pacific Biosciences (PacBio) HiFi long-read sequencing technology [81], which uses circular consensus sequencing to generate long-reads that are >99% accurate. The library preparation was performed using the SMRTbell® Express Template Prep Kit 2.0 following the manufacturer’s instructions (Pacific Biosciences). Additional information on the TELL-Seq™, stLFR, Hi-C and PacBio HiFi library preparations can be found in the electronic supplementary material.

### (c) Genome assembly and annotation

Different genome assembly pipelines were utilized for the different species depending on the sequencing resources available (Figure S1). *De novo* assemblies based on linked-read data were performed using the Supernova assembler v2.1.1 [82]. Prior to running Supernova, both the TELL-Seq and stLFR data were converted to a ‘10X format’: TELL-Seq sequences were converted using the ‘ust10x’ tool provided by Universal Sequencing Technology (UST), and the Supernova barcode whitelist was replaced by a TELL-Seq barcode whitelist consisting of ∼24.58 million barcodes, and the stLFR sequences were converted using the stLFR2Supernova pipeline (https://github.com/BGI-Qingdao/stlfr2supernova_pipeline) with the parameter BARCODE_FREQ_THRESHOLD set to ‘1’.

For the genome of *P. cribrata*, we performed a *de novo* assembly based on the PacBio Hifi sequencing data using Improved Phased Assembler (IPA) v1.3.1 (Pacific Biosciences, 2022) with default parameters. We then used *quickmerge* [83] to merge this IPA long-read *de novo* assembly (i.e., the ‘reference assembly’) to the Supernova linked-read *de novo* assembly (i.e., the ‘query assembly’), implementing default settings.

For the genome of *G. dilatatus*, gaps in the Supernova *de novo* assembly were filled using the program TGS-GapCloser v1.1.1 [84] with low-coverage PacBio Hifi data, implementing the ‘g_check’ flag (i.e., gap size checking for gaps >100 bp), and setting the minimap2 argument as ‘-x asm20’. The resultant genome assembly was then scaffolded using the HiC sequence data. First, Hi-C reads were mapped to the draft assembly using the pipeline provided by Arima Genomics (https://github.com/ArimaGenomics/mapping_pipeline). Second, the assembly was scaffolded based on the aligned Hi-C reads using SALSA2 [85,86], implementing the ‘CLEAN’ flag to check for misassemblies.

All three genome assemblies were annotated using FGENESH++ v7.2.2 [87,88], carried out on the Nimbus cloud service (provided by the Pawsey Supercomputing Centre). To prepare the assemblies for annotation, we generated a repeat database using RepeatModeler v2.0.1 [89], then masked repetitive regions in the assemblies using RepeatMasker v4.0.6 [90], implementing the ‘-nolow’ option (i.e., low complexity or simple repeat regions are not masked). Both masked and unmasked genome assemblies were used as input for FGENESH++. FGENESH++ was run with the non-mammalian general pipeline parameters, optimized gene-finding parameters trained on the genome of the pea aphid (*Acyrthosiphon pisum*), and a curated insect protein database provided by Softberry (2020 release) for the “prot_map” homology-based predictions. *Ab initio* predictions without any BLAST hits were removed. As an extra quality control step, we performed a protein-protein BLAST search of all annotated genes against the aforementioned insect protein database and investigated the sequence similarities and alignment lengths of the top hits. We generated various statistics to assess assembly continuity using QUAST v4.3 [91], and assessed assembly completeness using BUSCO v4.0.6 (utilizing the Insecta lineage data set [92–95]).

### (d) Orthology assignment

Proteomes of these three newly generated blaberid annotated genomes were combined with proteomes from six other species extracted from InsectBase 2.0 [96], including: American cockroach (*Periplaneta americana*), German cockroach (*Blatella germanica*), Nevada termite (*Zootermopsis nevadensis*), Formosan termite (*Coptotermes formosanus*), a drywood termite (*Cryptotermes secundus*), and a cricket (*Laupala kohalensis*) as an outgroup (Table S2). The cricket proteome was utilized as an outgroup rather than a proteome from a more closely related species (e.g., *Clitarchus hookeri*) due to its relatively high genome completeness. The longest isoform for each gene was extracted using the primary_transcript.py script supplied by OrthoFinder v2.4 [97].

Orthologous protein sequences among all nine species were allocated using OrthoFinder with default settings. Resultant orthogroups were annotated based on the most frequent gene annotation within the orthogroup. Single-copy orthologue sequences were identified and aligned using OrthoFinder, implementing the MAFFT sequence alignment method [98]. The corresponding nucleotide coding sequences (CDS) of these single-copy orthologue protein sequences were extracted from each of the respective genomes, and aligned using the codon-aware sequence alignment supplied by HyPhy v2.5.32 [28,29]. These single-copy orthologue CDS were utilized for our subsequent analyses of selection.

### (e) Gene expansions and contractions

The gene counts generated using OrthoFinder were used to investigate gene family expansions and contractions under a birth-death model using CAFE v.5.0 [99]. To prepare the input tree for CAFE, the rooted species tree generated by OrthoFinder was converted into an ultrametric tree using the make_ultrametric.py script supplied by OrthoFinder. Gene families with ≥100 genes in any one lineage were extracted using a CAFE python script (clade_and_size_filter.py) and analysed separately, because high variance of gene copy number can lead to noninformative parameter estimates. The orthogroup data sets were analysed with CAFE, implementing a Poisson root frequency distribution (-p), a model to account for genome assembly and annotation errors (-e), and a lambda tree (-y) specifying three separate lambda estimations for the termites, cockroaches, and outgroup (following results from CODEML analyses).

### (f) Positive and relaxed selection

We estimated *d_N_/d_S_* values for the aforementioned single-copy orthologues extracted from the nine analysed genomes using CODEML, which is part of the PAML package [26]. We used a fixed tree topology, based on previous phylogenetic estimates (Figure 2b [19,20,100]), and implemented various *d_N_/d_S_*partitions across the tree in separate analyses. First, we implemented the free-ratio model, which estimates a separate *d_N_* and *d_S_* value for each branch across the phylogeny. We ran CODEML, implementing the free-ratio model, three times on 1484 single-copy orthologue alignments (seven alignments were removed due to missingness), then extracted the estimated *d_N_* and *d_S_*values from the run with the highest likelihood score. To calculate the *d_N_*and *d_S_* rates, we scaled these values with branch lengths from chronograms derived from Evangelista et al. [20] and Beasley-Hall et al. [100].

Second, we implemented two- and three-ratio *d_N_/d_S_*models, whereby specific sets of branches are nominated as foreground branches (i.e., the branches ‘of interest’), and other branches are nominated as background branches. In the two-ratio model, we set the branch to the outgroup (*Laupala kohalensis*) as the background branch, allowing a single *d_N_/d_S_* value to be estimated across Blattodea. In the three-ratio model, we allowed separate *d_N_/d_S_*values to be estimated for the termite clade (excluding the termite stem branch) and the cockroach branches (i.e., the rest of Blattodea). We ran CODEML, implementing the two- and three-ratio models, for all 1484 single-copy orthologue alignments. For each orthologue, we used likelihood-ratio tests to evaluate whether separate values for *d_N_/d_S_* for the termites and other taxa in Blattodea fit the sequence data better than a single *d_N_/d_S_* value for Blattodea.

We investigated positive selection for each gene on different branches using an adaptive branch-site random-effects model, aBSREL [101], implemented in HyPhy. aBSREL estimates *d_N_/d_S_*values and tests whether a proportion of sites have evolved under positive selection for each user-specified test branch in the tree. First, we analysed the four termite branches (i.e., assigned them as the ‘foreground’ branches) to investigate signatures of positive selection related to the evolution of termites. Second, we analysed a larger subset of branches across the cockroach and termite phylogeny (i.e., 12 branches assigned as ‘foreground’ branches) to investigate broader patterns of positive selection in the group. Resultant *p*-values were corrected for multiple branches being tested using the Holm-Bonferroni method (based on the 5% level of significance). We repeated each analysis using three models accounting for multinucleotide mutations: multiple-hits ‘Double’, multiple hits ‘Double+Triple’, and the standard model. Analyses implementing the best-fitting model for each orthologue were retained.

To test for genes with evidence of relaxed selection, we applied the RELAX method [27], implemented in HyPhy [28,29], to each alignment separately. RELAX fits three *d_N_/d_S_*classes to the phylogeny then tests for relaxed/intensified selection on a user-specified test branch. We ran RELAX for 12 different test branches in separate analyses (Figure 1c). Resultant *p*-values were corrected for multiple branches being tested using the Holm-Bonferroni method (based on the 5% level of significance). To evaluate whether patterns of relaxed selection were due to errors in our automated sequence alignments, we repeated the RELAX analysis on a subset of audited orthologue alignments. We manually inspected and aligned sequences of 105 orthologues, including 20 randomly selected orthologues that were found to be under relaxed selection in all three termite species, 22 that were found to be under relaxed selection in two termite species, and 65 that yielded no evidence of relaxed selection. We subsequently carried out a RELAX analysis on the realigned orthologues, implementing the standard model for multinucleotide mutations.

### (g) Analysis of differentially expressed genes

We identified and analysed caste-biased and unbiased genes characterized by Harrison et al. [6] and Maekawa et al. [30] in our single-copy orthologue data set based on the putative annotations. We tested whether there were significantly more or fewer caste-biased orthogroups (and unbiased orthogroups) under relaxed or positive selection compared with the whole data set using chi-squared tests, implemented in the R package ‘stats’ [102].

To investigate the evolutionary history of these caste-biased orthogroups (and unbiased orthogroups), we assessed if they were associated with a higher or lower rate of evolution in cockroaches (see [8]). We did this by estimating total tree lengths for the cockroaches. First, we estimated phylograms for each single-copy orthologue using IQ-TREE v2.2.2 [103] using a fixed topology based on previous phylogenetic estimates as above (see Figure 2b), and applying ModelFinder [104] (implemented in IQ-TREE 2) to utilize the best-fitting substitution model for each orthologue. Second, all branch lengths, excluding the outgroup branch, were summed to obtain a total tree length for each orthologue. Third, all non-termite branches, including the cockroach terminal branches and associated internal branches, were summed to obtain a ‘cockroach’ tree length.

Fourth, this ‘cockroach’ tree length was divided by the total tree length to determine the relative cockroach tree length, or the relative contribution of cockroach (i.e., non-termite) substitutions to the evolution of the single-copy orthologues. Lastly, we investigated if there was an association between cockroach tree length and caste-biased genes via comparisons to the entire data set using a Mann-Whitney U test, implemented in the R package ‘stats.’ In addition to the tree-length analysis, we investigated the selection signatures of these caste-biased orthogroups in cockroaches, and compared them with the whole data set using chi-squared tests as above.

#### (h) Measuring the ‘genetic load’

To investigate genetic load among the sampled taxa, we analysed amino acid substitutions in BUSCOs from the Insecta lineage data set. These highly conserved genes are expected to be under purifying selection, and most derived alleles are likely to be deleterious [105,106]. Here, we focus on ‘realized load’ and not ‘masked load’, as heterozygosity was not considered [49,51].

For this analysis, we extracted BUSCOs from the Insecta lineage data set in each of the sampled taxa using BUSCO v5.4.2. It should be noted that the FGENESH++ pipeline that was used to annotate the three genomes produced in this study was developed to annotate thousands of genes with broad parameters, hence not all BUSCOs present in the genome assembly were annotated; consequently, some of the BUSCOs analysed (which were identified using a BUSCO bioinformatic pipeline; see Methods) were not present in the aforementioned single-copy orthologue data set. We only retained BUSCOs that were present in the genomes of all nine examined taxa, resulting in a total of 581 genes for this genetic load analysis.

We then polarized the amino acid substitutions of the aligned BUSCOs relative to the corresponding BUSCOs from the cricket outgroup. Each of the cockroach and termite BUSCOs were aligned with the cricket BUSCO using the codon-aware sequence alignment supplied by HyPhy. Gaps and ambiguities were masked, and SNPs were called with SNP-sites [107]. We then analysed these derived amino acid substitutions to infer realized genetic load based on three metrics: 1) the total number of derived substitutions per amino acid; 2) the number of putatively deleterious substitutions per amino acid; and 3) the ratio of putatively deleterious to benign substitutions. We used PROVEAN v1.1.5 [50] to predict the deleterious effect of derived substitutions based on how phylogenetically conserved the allele is amongst a database of homologous protein sequences [108–110]. The derived alleles were classified as either “benign” (PROVEAN scores > −2.5) or “deleterious” (PROVEAN scores ≤ −2.5). The total number of derived substitutions per amino acid was also included as a proxy for genetic load, as a previous study showed that this is a good predictor of fitness, and superior to utilizing PROVEAN in some cases [111]. The genetic load metrics of each BUSCO locus were compared between species using a paired Wilcoxon signed-rank test, implemented in the R package ‘stats’.

## Acknowledgements

We utilized the Artemis HPC cluster provided by the University of Sydney, the FlashLite HPC cluster provided by the University of Queensland’s Research Computing Centre (RCC), and the Nimbus cloud service provided by the Pawsey Supercomputing Centre. We thank Carolyn Hogg for advice on high performance computing resources.

## Data, code and materials

The genome assemblies and annotation will be made publicly available upon acceptance.

## Author contributions

K.M.E.: conceptualization, data curation, formal analysis, methodology, writing—original draft, writing—review and editing; S.Y.W.H.: conceptualization, methodology, funding acquisition, writing—review and editing; A-A.C.: formal analysis, methodology, writing— review and editing; F.R.J.: formal analysis, methodology, writing—review and editing; Y.K.: resources, data curation, writing—review and editing; J.B.: formal analysis, writing—review and editing; T.B.: resources, data curation, writing—review and editing; H.A.R.: resources, data curation, writing—review and editing; N.L.: conceptualization, supervision, methodology, resources, project administration, funding acquisition, writing—review and editing.

## References

1. Wilson, E. O. (1990). Success and dominance in ecosystems: The case of the social insects. Oldendorf, Germany: Ecology Institute.

2. Anderson, M. (1984). The evolution of eusociality. Annual Review of Ecology and Systematics, 15, 165–189.

3. Romiguier, J., Lourenco, J., Gayral, P., Faivre, N., Weinert, L. A., Ravel, S., … & Galtier, N. (2014). Population genomics of eusocial insects: the costs of a vertebrate-like effective population size. Journal of Evolutionary Biology, 27, 593–603.

4. Rubenstein, D. R., Ågren, J. A., Carbone, L., Elde, N. C., Hoekstra, H. E., Kapheim, K. M., … & Hofmann, H. A. (2019). Coevolution of genome architecture and social behavior. Trends in Ecology and Evolution, 34, 844–855.

5. Imrit, M. A., Dogantzis, K. A., Harpur, B. A., & Zayed, A. (2020). Eusociality influences the strength of negative selection on insect genomes. Proceedings of the Royal Society B, 287, 20201512.

6. Harrison, M. C., Chernyshova, A. M., & Thompson, G. J. (2021). No obvious transcriptome-wide signature of indirect selection in termites. Journal of Evolutionary Biology, 34, 403–415.

7. Hunt, B. G., Wyder, S., Elango, N., Werren, J. H., Zdobnov, E. M., Yi, S. V., & Goodisman, M. A. (2010). Sociality is linked to rates of protein evolution in a highly social insect. Molecular Biology and Evolution, 27, 497–500.

8. Hunt, B. G., Ometto, L., Wurm, Y., Shoemaker, D., Yi, S. V., Keller, L., & Goodisman, M. A. (2011). Relaxed selection is a precursor to the evolution of phenotypic plasticity. Proceedings of the National Academy of Sciences of the U.S.A., 108, 15936–15941.

9. Helanterä, H., & Uller, T. (2014). Neutral and adaptive explanations for an association between caste-biased gene expression and rate of sequence evolution. Frontiers in Genetics, 5, 297.

10. Settepani, V., Bechsgaard, J., & Bilde, T. (2016). Phylogenetic analysis suggests that sociality is associated with reduced effectiveness of selection. Ecology and Evolution, 6, 469–477.

11. Dyson, C. J., Piscano, O. L., Durham, R. M., Thompson, V. J., Johnson, C. H., & Goodisman, M. A. (2021). Temporal analysis of effective population size and mating system in a social wasp. Journal of Heredity, 112, 626–634.

12. Weyna, A., & Romiguier, J. (2021). Relaxation of purifying selection suggests low effective population size in eusocial Hymenoptera and solitary pollinating bees. Peer Community Journal, 1, e2.

13. Bromham, L., & Leys, R. (2005). Sociality and the rate of molecular evolution. Molecular Biology and Evolution, 22, 1393–1402.

14. Ellegren, H., & Parsch, J. (2007). The evolution of sex-biased genes and sex-biased gene expression. Nature Reviews Genetics, 8, 689–698.

15. Hall, D. W., & Goodisman, M. A. (2012). The effects of kin selection on rates of molecular evolution in social insects. Evolution, 66, 2080–2093.

16. Lo, N., Tokuda, G., Watanabe, H., Rose, H., Slaytor, M., Maekawa, K., … & Noda, H. (2000). Evidence from multiple gene sequences indicates that termites evolved from wood-feeding cockroaches. Current Biology, 10, 801–804.

17. Inward, D., Beccaloni, G., & Eggleton, P. (2007). Death of an order: a comprehensive molecular phylogenetic study confirms that termites are eusocial cockroaches. Biology Letters, 3, 331–335.

18. Legendre, F., Nel, A., Svenson, G. J., Robillard, T., Pellens, R., & Grandcolas, P. (2015). Phylogeny of Dictyoptera: dating the origin of cockroaches, praying mantises and termites with molecular data and controlled fossil evidence. PLOS ONE, 10, e0130127.

19. Bourguignon, T., Tang, Q., Ho, S. Y. W., Juna, F., Wang, Z., Arab, D. A., … & Lo, N. (2018). Transoceanic dispersal and plate tectonics shaped global cockroach distributions: evidence from mitochondrial phylogenomics. Molecular Biology and Evolution, 35, 970–983.

20. Evangelista, D. A., Wipfler, B., Béthoux, O., Donath, A., Fujita, M., Kohli, M. K., … & Simon, S. (2019). An integrative phylogenomic approach illuminates the evolutionary history of cockroaches and termites (Blattodea). Proceedings of the Royal Society B, 286, 20182076.

21. Terrapon, N., Li, C., Robertson, H. M., Ji, L., Meng, X., Booth, W., … & Liebig, J. (2014). Molecular traces of alternative social organization in a termite genome. Nature Communications, 5, 3636.

22. Harrison, M. C., Jongepier, E., Robertson, H. M., Arning, N., Bitard-Feildel, T., Chao, H., … & Bornberg-Bauer, E. (2018). Hemimetabolous genomes reveal molecular basis of termite eusociality. Nature Ecology and Evolution, 2, 557–566.

23. Itakura, S., Yoshikawa, Y., Togami, Y., & Umezawa, K. (2020). Draft genome sequence of the termite, *Coptotermes formosanus*: Genetic insights into the pyruvate dehydrogenase complex of the termite. Journal of Asia-Pacific Entomology, 23, 666–674.

24. Shigenobu, S., Hayashi, Y., Watanabe, D., Tokuda, G., Hojo, M. Y., Toga, K., … & Maekawa, K. (2022). Genomic and transcriptomic analyses of the subterranean termite *Reticulitermes speratus*: Gene duplication facilitates social evolution. Proceedings of the National Academy of Sciences of the U.S.A., 119, e2110361119.

25. Li, S., Zhu, S., Jia, Q., Yuan, D., Ren, C., Li, K., … & Zhan, S. (2018). The genomic and functional landscapes of developmental plasticity in the American cockroach. Nature Communications, 9, 1008.

26. Yang, Z. (2007). PAML 4: Phylogenetic analysis by maximum likelihood. Molecular Biology and Evolution, 24, 1586–1591.

27. Wertheim, J. O., Murrell, B., Smith, M. D., Kosakovsky Pond, S. L., & Scheffler, K. (2015). RELAX: detecting relaxed selection in a phylogenetic framework. Molecular Biology and Evolution, 32, 820–832.

28. Pond, S. L. K., Frost, S. D., & Muse, S. V. (2005). HyPhy: hypothesis testing using phylogenies. Bioinformatics, 21, 676–679.

29. Pond, S. L. K., Poon, A. F., Velazquez, R., Weaver, S., Hepler, N. L., Murrell, B., … & Muse, S. V. (2020). HyPhy 2.5—a customizable platform for evolutionary hypothesis testing using phylogenies. Molecular Biology and Evolution, 37, 295–299.

30. Maekawa, K., Hayashi, Y., & Lo, N. (2022). Termite sociogenomics: evolution and regulation of caste-specific expressed genes. Current Opinion in Insect Science, 50, 100880.

31. Wright, S. (1931). Evolution in mendelian populations. Genetics, 16, 97–159.

32. Nei, M., & Tajima, F. (1981). Genetic drift and estimation of effective population size. Genetics, 98, 625–640.

33. Crozier, R. H. (1979). Genetics of sociality. Social Insects, 1, 223–286.

34. Glémin, S. (2007). Mating systems and the efficacy of selection at the molecular level. Genetics, 177, 905–916.

35. Kimura, M. (1983). The Neutral Theory of Molecular Evolution. Cambridge, UK: Cambridge University Press.

36. Lynch, M., Ackerman, M. S., Gout, J. F., Long, H., Sung, W., Thomas, W. K., & Foster, P. L. (2016). Genetic drift, selection and the evolution of the mutation rate. Nature Reviews Genetics, 17, 704–714.

37. Chak, S. T., Baeza, J. A., & Barden, P. (2021). Eusociality shapes convergent patterns of molecular evolution across mitochondrial genomes of snapping shrimps. Molecular Biology and Evolution, 38, 1372–1383.

38. Bergeron, L. A., Besenbacher, S., Zheng, J., Li, P., Bertelsen, M. F., Quintard, B., … & Zhang, G. (2023). Evolution of the germline mutation rate across vertebrates. Nature, 615, 285–291.

39. Leichty, A. R., Pfennig, D. W., Jones, C. D., & Pfennig, K. S. (2012). Relaxed genetic constraint is ancestral to the evolution of phenotypic plasticity. Integrative and Comparative Biology, 52, 16–30.

40. Schrader, L., Helanterä, H., & Oettler, J. (2017). Accelerated evolution of developmentally biased genes in the tetraphenic ant *Cardiocondyla obscurior*. Molecular Biology and Evolution, 34, 535–544.

41. Traniello, J. F., Rosengaus, R. B., & Savoie, K. (2002). The development of immunity in a social insect: evidence for the group facilitation of disease resistance. Proceedings of the National Academy of Sciences of the U.S.A., 99, 6838–6842.

42. Cremer, S., Armitage, S. A., & Schmid-Hempel, P. (2007). Social immunity. Current Biology, 17, R693–R702.

43. Meusemann, K., Korb, J., Schughart, M., & Staubach, F. (2020). No evidence for single-copy immune-gene specific signals of selection in termites. Frontiers in Ecology and Evolution, 8, 26.

44. He, S., Sieksmeyer, T., Che, Y., Mora, M. A. E., Stiblik, P., Banasiak, R., … & McMahon, D. P. (2021). Evidence for reduced immune gene diversity and activity during the evolution of termites. Proceedings of the Royal Society B, 288, 20203168.

45. Bulmer, M. S., & Stefano, A. M. (2022). Termite eusociality and contrasting selective pressure on social and innate immunity. Behavioral Ecology and Sociobiology, 76, 4.

46. Pascoal, S., Shimadzu, H., Mashoodh, R., & Kilner, R. M. (2023). Parental care results in a greater mutation load, for which it is also a phenotypic antidote. Proceedings of the Royal Society B, 290, 20230115.

47. Tong, K. J., Duchêne, S., Ho, S. Y. W., & Lo, N. (2015). Comment on “Phylogenomics resolves the timing and pattern of insect evolution”. Science, 349, 487.

48. Muller, H. J. (1950). Our load of mutations. American Journal of Human Genetics, 2, 111–176.

49. Crow, J. F. (1970). Genetic loads and the cost of natural selection. In: Kojima, K.-i. (ed) Mathematical Topics in Population Genetics. Biomathematics, vol 1. Berlin, Germany: Springer.

50. Choi, Y., Sims, G. E., Murphy, S., Miller, J. R., & Chan, A. P. (2012). Predicting the functional effect of amino acid substitutions and indels. PLOS ONE, 7, e46688

51. Bertorelle, G., Raffini, F., Bosse, M., Bortoluzzi, C., Iannucci, A., Trucchi, E., … & Van Oosterhout, C. (2022). Genetic load: genomic estimates and applications in non-model animals. Nature Reviews Genetics, 23, 492–503.

52. Frankham, R., Bradshaw, C. J., & Brook, B. W. (2014). Genetics in conservation management: revised recommendations for the 50/500 rules, Red List criteria and population viability analyses. Biological Conservation, 170, 56–63.

53. Frankham, R., Ballou, J. D., Ralls, K., Eldridge, M., Dudash, M. R., Fenster, C. B., … & Sunnucks, P. (2017). Genetic management of fragmented animal and plant populations. Oxford, UK: Oxford University Press.

54. Thorne, B. L., Traniello, J. F. A., Adams, E. S., & Bulmer, M. (1999). Reproductive dynamics and colony structure of subterranean termites of the genus *Reticulitermes* (Isoptera Rhinotermitidae): a review of the evidence from behavioral, ecological, and genetic studies. Ethology Ecology & Evolution, 11, 149–169.

55. Lepage, M., & Darlington, J. P. (2000). Population dynamics of termites. In: Abe, T., Bignell, D.E., & Higashi, M. (eds) Termites: Evolution, Sociality, Symbioses, Ecology (pp. 333–361). Dordrecht, The Netherlands: Springer.

56. Smith, C. R., Toth, A. L., Suarez, A. V., & Robinson, G. E. (2008). Genetic and genomic analyses of the division of labour in insect societies. Nature Reviews Genetics, 9, 735–748.

57. DeHeer, C. J., & Vargo, E. L. (2006). An indirect test of inbreeding depression in the termites *Reticulitermes flavipes* and *Reticulitermes virginicus*. Behavioral Ecology and Sociobiology, 59, 753–761.

58. Rosengaus, R. B., & Traniello, J. F. (1993). Disease risk as a cost of outbreeding in the termite *Zootermopsis angusticollis*. Proceedings of the National Academy of Sciences of the U.S.A., 90, 6641–6645.

59. Fei, H. X., & Henderson, G. (2003). Comparative study of incipient colony development in the Formosan subterranean termite, *Coptotermes formosanus* Shiraki (Isoptera, Rhinotermitidae). Insectes Sociaux, 50, 226–233.

60. Calleri, D. V., Rosengaus, R. B., & Traniello, J. F. (2005). Disease and colony foundation in the dampwood termite *Zootermopsis angusticollis*: the survival advantage of nestmate pairs. Naturwissenschaften, 92, 300–304.

61. Eyer, P. A., & Vargo, E. L. (2022). Short and long-term costs of inbreeding in the lifelong-partnership in a termite. Communications Biology, 5, 389.

62. Lyko, F., Foret, S., Kucharski, R., Wolf, S., Falckenhayn, C., & Maleszka, R. (2010). The honey bee epigenomes: differential methylation of brain DNA in queens and workers. PLOS Biology, 8, e1000506.

63. Bonasio, R., Li, Q., Lian, J., Mutti, N. S., Jin, L., Zhao, H., … & Reinberg, D. (2012). Genome-wide and caste-specific DNA methylomes of the ants *Camponotus floridanus* and *Harpegnathos saltator*. Current Biology, 22, 1755–1764.

64. Kapheim, K. M., Pan, H., Li, C., Salzberg, S. L., Puiu, D., Magoc, T., … & Zhang, G. (2015). Genomic signatures of evolutionary transitions from solitary to group living. Science, 348, 1139–1143.

65. Glastad, K. M., Gokhale, K., Liebig, J., & Goodisman, M. A. (2016). The caste-and sex-specific DNA methylome of the termite *Zootermopsis nevadensis*. Scientific Reports, 6, 37110.

66. Oldroyd, B. P., & Yagound, B. (2021). The role of epigenetics, particularly DNA methylation, in the evolution of caste in insect societies. Philosophical Transactions of the Royal Society B, 376, 20200115.

67. Sieber, K. R., Dorman, T., Newell, N., & Yan, H. (2021). (Epi)genetic mechanisms underlying the evolutionary success of eusocial insects. Insects, 12, 498.

68. Cedar, H., & Bergman, Y. (2009). Linking DNA methylation and histone modification: patterns and paradigms. Nature Reviews Genetics, 10, 295–304.

69. Vaughan, R. M., Kupai, A., & Rothbart, S. B. (2021). Chromatin regulation through ubiquitin and ubiquitin-like histone modifications. Trends in Biochemical Sciences, 46, 258–269.

70. Spannhoff, A., Kim, Y. K., Raynal, N. J. M., Gharibyan, V., Su, M. B., Zhou, Y. Y., … & Bedford, M. T. (2011). Histone deacetylase inhibitor activity in royal jelly might facilitate caste switching in bees. EMBO Reports, 12, 238–243.

71. Dickman, M. J., Kucharski, R., Maleszka, R., & Hurd, P. J. (2013). Extensive histone post-translational modification in honey bees. Insect Biochemistry and Molecular Biology, 43, 125–137.

72. Wojciechowski, M., Lowe, R., Maleszka, J., Conn, D., Maleszka, R., & Hurd, P. J. (2018). Phenotypically distinct female castes in honey bees are defined by alternative chromatin states during larval development. Genome Research, 28, 1532–1542.

73. Simola, D. F., Ye, C., Mutti, N. S., Dolezal, K., Bonasio, R., Liebig, J., … & Berger, S. L. (2013). A chromatin link to caste identity in the carpenter ant *Camponotus floridanus*. Genome Research, 23, 486–496.

74. Simola, D. F., Graham, R. J., Brady, C. M., Enzmann, B. L., Desplan, C., Ray, A., … & Berger, S. L. (2016). Epigenetic (re) programming of caste-specific behavior in the ant *Camponotus floridanus*. Science, 351, aac6633.

75. Warner, M. R., Qiu, L., Holmes, M. J., Mikheyev, A. S., & Linksvayer, T. A. (2019). Convergent eusocial evolution is based on a shared reproductive groundplan plus lineage-specific plastic genes. Nature Communications, 10, 2651.

76. Wang, J., Santiago, E., & Caballero, A. (2016). Prediction and estimation of effective population size. Heredity, 117, 193–206.

77. Chen, Z., Pham, L., Wu, T. C., Mo, G., Xia, Y., Chang, P. L., … & Lei, M. (2020). Ultralow-input single-tube linked-read library method enables short-read second-generation sequencing systems to routinely generate highly accurate and economical long-range sequencing information. Genome Research, 30, 898–909.

78. Wang, O., Chin, R., Cheng, X., Wu, M. K. Y., Mao, Q., Tang, J., … & Peters, B. A. (2019). Efficient and unique cobarcoding of second-generation sequencing reads from long DNA molecules enabling cost-effective and accurate sequencing, haplotyping, and de novo assembly. Genome Research, 29, 798–808.

79. Belton, J. M., McCord, R. P., Gibcus, J. H., Naumova, N., Zhan, Y., & Dekker, J. (2012). Hi–C: a comprehensive technique to capture the conformation of genomes. Methods, 58, 268–276.

80. DeMaere, M. Z., & Darling, A. E. (2021). qc3C: reference-free quality control for Hi-C sequencing data. PLOS Computational Biology, 17, e1008839.

81. Wenger, A. M., Peluso, P., Rowell, W. J., Chang, P. C., Hall, R. J., Concepcion, G. T., … & Hunkapiller, M. W. (2019). Accurate circular consensus long-read sequencing improves variant detection and assembly of a human genome. Nature Biotechnology, 37, 1155–1162.

82. Weisenfeld, N. I., Kumar, V., Shah, P., Church, D. M., & Jaffe, D. B. (2017). Direct determination of diploid genome sequences. Genome Research, 27, 757–767.

83. Chakraborty, M., Baldwin-Brown, J. G., Long, A. D., & Emerson, J. J. (2016). Contiguous and accurate de novo assembly of metazoan genomes with modest long read coverage. Nucleic Acids Research, 44, e147.

84. Xu, M., Guo, L., Gu, S., Wang, O., Zhang, R., Peters, B. A., … & Zhang, Y. (2020). TGS-GapCloser: a fast and accurate gap closer for large genomes with low coverage of error-prone long reads. GigaScience, 9, giaa094.

85. Ghurye, J., Pop, M., Koren, S., Bickhart, D., & Chin, C. S. (2017). Scaffolding of long read assemblies using long range contact information. BMC Genomics, 18, 527.

86. Ghurye, J., Rhie, A., Walenz, B. P., Schmitt, A., Selvaraj, S., Pop, M., … & Koren, S. (2019). Integrating Hi-C links with assembly graphs for chromosome-scale assembly. PLOS Computational Biology, 15, e1007273.

87. Salamov, A.A. & Solovyev, V.V. (2000) Ab initio gene finding in *Drosophila* genomic DNA. Genome Research, 10, 516–522.

88. Solovyev, V., Kosarev, P., Seledsov, I., & Vorobyev, D. (2006). Automatic annotation of eukaryotic genes, pseudogenes and promoters. Genome Biology, 7, S10.

89. Flynn, J. M., Hubley, R., Goubert, C., Rosen, J., Clark, A. G., Feschotte, C., & Smit, A. F. (2020). RepeatModeler2 for automated genomic discovery of transposable element families. Proceedings of the National Academy of Sciences of the U.S.A., 117, 9451–9457.

90. Smit, A., Hubley, R., & Green, P. (2015). RepeatMasker Open-4.0. Retrieved from http://www.repeatmasker.org.

91. Gurevich, A., Saveliev, V., Vyahhi, N., & Tesler, G. (2013). QUAST: quality assessment tool for genome assemblies. Bioinformatics, 29, 1072–1075.

92. Simão, F. A., Waterhouse, R. M., Ioannidis, P., Kriventseva, E. V., & Zdobnov, E. M. (2015). BUSCO: assessing genome assembly and annotation completeness with single-copy orthologs. Bioinformatics, 31, 3210–3212.

93. Waterhouse, R. M., Seppey, M., Simão, F. A., Manni, M., Ioannidis, P., Klioutchnikov, G., … & Zdobnov, E. M. (2018). BUSCO applications from quality assessments to gene prediction and phylogenomics. Molecular Biology and Evolution, 35, 543–548.

94. Waterhouse, R. M., Seppey, M., Simão, F. A., & Zdobnov, E. M. (2019). Using BUSCO to assess insect genomic resources. Methods in Molecular Biology, 1858, 59– 74.

95. Manni, M., Berkeley, M. R., Seppey, M., & Zdobnov, E. M. (2021). BUSCO: assessing genomic data quality and beyond. Current Protocols, 1, e323.

96. Mei, Y., Jing, D., Tang, S., Chen, X., Chen, H., Duanmu, H., … & Li, F. (2022). InsectBase 2.0: a comprehensive gene resource for insects. Nucleic Acids Research, 50, D1040–D1045.

97. Emms, D. M., & Kelly, S. (2019). OrthoFinder: phylogenetic orthology inference for comparative genomics. Genome Biology, 20, 238.

98. Katoh, K., & Standley, D. M. (2013). MAFFT multiple sequence alignment software version 7: improvements in performance and usability. Molecular Biology and Evolution, 30, 772–780.

99. Mendes, F.K., Vanderpool, D., Fulton, B., & Hahn, M.W. (2020) CAFÉ 5 models variation in evolutionary rates among gene families. Bioinformatics, 36, 5516–5518.

100. Beasley-Hall, P. G., Rose, H. A., Walker, J., Kinjo, Y., Bourguignon, T., & Lo, N. (2021). Digging deep: a revised phylogeny of Australian burrowing cockroaches (Blaberidae: Panesthiinae, Geoscapheinae) confirms extensive nonmonophyly and provides insights into biogeography and evolution of burrowing. Systematic Entomology, 46, 767–783.

101. Smith, M. D., Wertheim, J. O., Weaver, S., Murrell, B., Scheffler, K., & Kosakovsky Pond, S. L. (2015). Less is more: an adaptive branch-site random effects model for efficient detection of episodic diversifying selection. Molecular Biology and Evolution, 32, 1342–1353.

102. R Core Team (2022). R: A language and environment for statistical computing. R Foundation for Statistical Computing, Vienna, Austria. URL https://www.R-project.org/.

103. Minh, B. Q., Schmidt, H. A., Chernomor, O., Schrempf, D., Woodhams, M. D., Von Haeseler, A., & Lanfear, R. (2020). IQ-TREE 2: new models and efficient methods for phylogenetic inference in the genomic era. Molecular Biology and Evolution, 37, 1530– 1534.

104. Kalyaanamoorthy, S., Minh, B. Q., Wong, T. K., Von Haeseler, A., & Jermiin, L. S. (2017). ModelFinder: fast model selection for accurate phylogenetic estimates. Nature Methods, 14, 587–589.

105. Eyre-Walker, A., & Keightley, P. D. (2007). The distribution of fitness effects of new mutations. Nature Reviews Genetics, 8, 610–618.

106. Boyko, A. R., Williamson, S. H., Indap, A. R., Degenhardt, J. D., Hernandez, R. D., Lohmueller, K. E., … & Bustamante, C. D. (2008). Assessing the evolutionary impact of amino acid mutations in the human genome. PLOS Genetics, 4, e1000083.

107. Page, A. J., Taylor, B., Delaney, A. J., Soares, J., Seemann, T., Keane, J. A., & Harris, S. R. (2016). SNP-sites: rapid efficient extraction of SNPs from multi-FASTA alignments. Microbial Genomics, 2, e000056.

108. Perrier, C., Ferchaud, A. L., Sirois, P., Thibault, I., & Bernatchez, L. (2017). Do genetic drift and accumulation of deleterious mutations preclude adaptation? Empirical investigation using RAD seq in a northern lacustrine fish. Molecular Ecology, 26, 6317– 6335.

109. Ochoa, A., & Gibbs, H. L. (2021). Genomic signatures of inbreeding and mutation load in a threatened rattlesnake. Molecular Ecology, 30, 5454–5469.

110. Kleinman-Ruiz, D., Lucena-Perez, M., Villanueva, B., Fernández, J., Saveljev, A. P., Ratkiewicz, M., … & Godoy, J. A. (2022). Purging of deleterious burden in the endangered Iberian lynx. Proceedings of the National Academy of Sciences of the U.S.A., 119, e2110614119.

111. Sandell, L., & Sharp, N. P. (2022). Fitness effects of mutations: An assessment of PROVEAN predictions using mutation accumulation data. Genome Biology and Evolution, 14, evac004.

