## Supplementary data for "Pervasive relaxed selection in termite genomes"

**Supplementary Material**

**Supplemental Tables**

**Table S1.** Types of DNA sequencing produced for the three cockroach genomes.

| **Species** | **Tell-Seq linked reads** | **stLFR linked reads** | **HiC** | **Pacbio HiFi** |
| --- | --- | --- | --- | --- |
| *Geoscapheus dilatatus* |  | **X** | **X** | **X** |
| *Neogeoscapheus hanni* |  | **X** |  |  |
| *Panesthia cribrata* | **X** |  |  | **X** |

**Table S2.** Genome statistics for the six proteomes utilized in the comparative genomic analyses. These statistics were retrieved from InsectBase2.0: <http://v2.insect-genome.com/>.

| **Species** | **Genome length** | **N50** | **Complete BUSCOs** | **Fragmented BUSCOs** | **Missing BUSCOs** | **Duplicated BUSCOs** |
| --- | --- | --- | --- | --- | --- | --- |
| *Blattella germanica* | 2.06 Gb | 1.07 Mb | 96.1% | 1.7% | 2.2% | 2.2% |
| *Periplaneta americana* | 3.42 Gb | 337 kb | 91.9% | 4.8% | 3.3% | 5.3% |
| *Cryptotermes secundus* | 1.03 Gb | 1.20 Mb | 97.4% | 1.0% | 1.6% | 1.4% |
| *Coptotermes formosanus* | 887 Mb | 1.44 Mb | 98.2% | 1.0% | 0.8% | 1.6% |
| *Zootermopsis nevadensis* | 491 Mb | 761 kb | 98.4% | 0.9% | 0.7% | 0.5% |
| *Laupala kohalensis* | 1.60 Gb | 584 kb | 97.9% | 0.9% | 1.2% | 1.8% |

**Table S3.** Proportion of audited orthogroups under relaxed selection on each terminal branch based on RELAX analyses (after a Holm-Bonferonni correction for testing multiple branches). The 105 audited orthologues (i.e., audited via manual alignments) comprise 20 orthologues that were found to be under relaxed selection in all 3 termite species in the original analyses (i.e., automated alignments), 22 that were found to be under relaxed selection in 2 termite species, and 65 where there was no evidence of relaxed selection.

| **Terminal branch** | **Proportion of orthologues (*n* = 105) under relaxed selection** | |
| --- | --- | --- |
|  | *Original alignment* | *Audited alignment* |
| *Cryptotermes secundus* | 30.48% | 26.67% |
| *Coptotermes formosanus* | 38.10% | 31.43% |
| *Zootermopsis nevadensis* | 30.48% | 34.29% |
| Cockroach species (*n* = 5) | 1.90–7.62% | 0–5.71% |

**Table S4.** Selection signature for caste-bias genes from Harrison et al. (2021) and Maekawa et al. (2022) that were identified in our data set. *Coptotermes formosanus* was not included as this species was not analysed by Harrison et al. (2021) or Maekawa et al. (2022). The ‘unbiased’ genes were only considered if they were unbiased in both *Zootermopsis nevadensis* and *Cryptotermes secundus*. There was no significant difference between the proportion of all single-copy orthologues (*n* = 1490) that were under relaxed or positive selection and unbiased or caste-bias orthogroups, including when worker- and queen-biased genes were combined; however, sample sizes for the worker-biased genes were too small to perform chi-square testing. If indirect selection was driving patterns of relaxed selection, the expectation is a greater proportion of caste-biased genes under relaxed selection (compared with the full data set), and relatively fewer unbiased genes under relaxed selection. Orthogroups under positive selection were based on the analysis in which foreground branches were assigned to termite taxa.

| **Data set** | **Orthogroups under relaxed selection in *Zootermopsis nevadensis*** | **Orthogroups under relaxed selection in *Cryptotermes secundus*** | **Orthogroups under positive selection in *Zootermopsis nevadensis*** | **Orthogroups under positive selection in *Cryptotermes secundus*** |
| --- | --- | --- | --- | --- |
| All orthogroups | 28.19% (420/1490) | 24.09% (359/1490) | 4.56% (68/1491) | 4.56% (68/1491) |
| Unbiased | 25.33% (38/150) | 20.00% (30/150) | 4.00% (6/150) | 4.67% (7/150) |
| Worker-biased in *Zootermopsis nevadensis* | 37.5% (3/8) | NA | 0% (0/8) | NA |
| Worker-biased in *Cryptotermes secundus* | NA | 0% (0/3) | NA | 0% (0/3) |
| Queen-biased in *Zootermopsis nevadensis* | 33.02% (35/106) | NA | 3.77% (4/106) | NA |
| Queen-biased in *Cryptotermes secundus* | NA | 25.30% (21/83) | NA | 1.20% (1/83) |

**Table S5.** Pairwise comparisons of genetic load metrics, including the number of substitutions per amino acid (‘subs/AA’), the number of putative deleterious substitutions (inferred using PROVEAN) per amino acid (‘del/AA’), and the ratio of putative deleterious to putative benign substitutions (‘del/ben’).

N = no significant difference.

**H** = the value associated with the row taxon (the vertical list) is significantly higher than the value associated with the column taxon.

**L** = the value associated with the row taxon is significantly lower than the value associated with the column taxon.

|  | ***C. formasanus*** | | | ***C. secundus*** | | | ***Z. nevadensis*** | | | ***P. americana*** | | | ***B. germanica*** | | | ***G. dilatatus*** | | | ***N. hanni*** | | |
| --- | --- | --- | --- | --- | --- | --- | --- | --- | --- | --- | --- | --- | --- | --- | --- | --- | --- | --- | --- | --- | --- |
|  | subs/AA | del/AA | del/ben | subs/AA | del/AA | del/ben | subs/AA | del/AA | del/ben | subs/AA | del/AA | del/ben | subs/AA | del/AA | del/ben | subs/AA | del/AA | del/ben | subs/AA | del/AA | del/ben |
| ***C. secundus*** | N | N | N |  |  |  |  |  |  |  |  |  |  |  |  |  |  |  |  |  |  |
| ***Z. nevadensis*** | N | N | N | N | N | N |  |  |  |  |  |  |  |  |  |  |  |  |  |  |  |
| ***P. americana*** | **L** | **H** | **H** | **L** | N | **H** | **L** | **H** | **H** |  |  |  |  |  |  |  |  |  |  |  |  |
| ***B. germanica*** | N | N | N | N | N | N | **L** | N | **H** | **H** | N | **L** |  |  |  |  |  |  |  |  |  |
| ***G. dilatatus*** | **L** | N | **H** | N | N | **H** | **L** | **H** | **H** | **H** | N | **L** | N | N | N |  |  |  |  |  |  |
| ***N. hanni*** | **L** | N | **H** | **L** | N | **H** | **L** | N | **H** | **H** | N | N | N | N | N | N | N | N |  |  |  |
| ***P. cribrata*** | **L** | N | **H** | **L** | N | **H** | **L** | N | **H** | **H** | N | **L** | N | N | N | **L** | N | N | N | N | N |

**Table S6.** List of orthogroups that had significant expansions/contractions on least one termite branches (*p* < 4×10^-5^) (when there was either an expansion or contraction on all significant branches). Orthogroups with uncharacterized annotations were excluded.

| **Orthogroup** | **Putative annotation** | **Contraction/**  **Expansion** | **Significant branches** |
| --- | --- | --- | --- |
| OG0000041 | Tubulin alpha-1 chain (alphaTub84B) | Contraction | *Coptotermes formosanus* terminal branch |
| OG0000046 | Lipase 3 (Lip3) | Contraction | *Coptotermes formosanus* terminal branch |
| OG0000050 | Craniofacial development protein 2 (CFDP2) | Expansion | *Coptotermes formosanus* terminal branch |
| OG0000075 | Probable RNA-directed DNA polymerase from transposon BS (RTase) | Expansion | *Coptotermes formosanus* terminal branch |
| OG0000114 | UDP-glycosyltransferase UGT5 (UGT5) | Expansion | *Coptotermes formosanus* terminal branch |
| OG0000154 | Tubulin beta-1 chain (TUBB) | Expansion | *Cryptotermes secundus* terminal branch |
| OG0000204 | Esterase FE4 | Expansion | *Cryptotermes secundus* terminal branch |
| OG0000238 | Nose resistant to fluoxetine protein 6 (nrf-6) | Expansion | *Cryptotermes secundus* terminal branch |
| OG0000343 | Fizzy-related protein homolog (Fzr1) | Contraction | *Coptotermes formosanus* terminal branch |
| OG0000425 | Lysozyme c-1 (Lyz1) | Expansion | *Cryptotermes secundus* terminal branch |
| OG0000453 | Protein argonaute-2 (Ago2) | Expansion | *Cryptotermes secundus* terminal branch |
| OG0000462 | Cuticle protein 2 | Contraction | *Coptotermes formosanus* terminal branch |
| OG0000492 | KAT8 regulatory NSL complex subunit 3 (Kansl3) | Expansion | *Coptotermes formosanus* terminal branch |
| OG0000493 | Serine/threonine-protein phosphatase 2A 56 kDa regulatory subunit epsilon isoform (PPP2R5E) | Expansion | *Cryptotermes secundus* terminal branch |
| OG0000581 | Geranylgeranyl pyrophosphate synthase (GGPS1) | Expansion | *Coptotermes formosanus* terminal branch |
| OG0000584 | Histone H4 transcription factor (HINFP) | Expansion | *Cryptotermes secundus* terminal branch |
| OG0000732 | Cytochrome P450 9e2 (CYP9E2) | Expansion | *Coptotermes formosanus* terminal branch |
| OG0000976 | E3 ubiquitin-protein ligase (Siah1a) | Expansion | *Cryptotermes secundus* terminal branch |
| OG0001006 | DNA/RNA-binding protein KIN17 (Kin) | Expansion | *Cryptotermes secundus* terminal branch |
| OG0001144 | Sphingomyelin phosphodiesterase (SMPD1) | Contraction | *Coptotermes formosanus* terminal branch |
| OG0001149 | Ubiquitin carboxyl-terminal hydrolase 45 (Usp45) | Expansion | *Zootermopsis nevadensis* terminal branch |
| OG0001322 | Protein roadkill (rdx) | Expansion | *Cryptotermes secundus* terminal branch |
| OG0001354 | DNA-binding protein RFX2 (Rfx2) | Expansion | *Cryptotermes secundus* terminal branch |
| OG0001362 | Kinesin-like protein (Klp10A) | Expansion | *Cryptotermes secundus* terminal branch |
| OG0001412 | Ribosomal protein S6 kinase alpha-5 (RPS6KA5) | Expansion | *Cryptotermes secundus* terminal branch |
| OG0001597 | Protein transport protein (Sec24C) | Expansion | *Zootermopsis nevadensis* terminal branch |
| OG0001661 | Pre-mRNA cleavage complex 2 protein (Pcf11) | Expansion | *Cryptotermes secundus* terminal branch |
| OG0001800 | Embryonic polarity protein dorsal (dl) | Expansion | *Zootermopsis nevadensis* terminal branch |
| OG0001844 | Forkhead transcription factor (HCM1) | Expansion | *Zootermopsis nevadensis* terminal branch |
| OG0001939 | Membrane alanyl aminopeptidase (ANPEP) | Expansion | *Coptotermes formosanus* terminal branch |
| OG0002110 | Angio-associated migratory cell protein (AAMP) | Expansion | *Zootermopsis nevadensis* terminal branch |
| OG0002126 | T-cell activation inhibitor, mitochondrial (TCAIM) | Expansion | *Coptotermes formosanus* terminal branch |
| OG0002278 | Cell division cycle 5-like protein (CDC5L) | Expansion | *Cryptotermes secundus* terminal branch |
| OG0002282 | Cell division cycle protein 20 homolog (Cdc20) | Expansion | *Zootermopsis nevadensis* terminal branch |
| OG0002314 | Rab3 GTPase-activating protein non-catalytic subunit (Rab3gap2) | Expansion | *Zootermopsis nevadensis* terminal branch |
| OG0002315 | Protein brambleberry (bmb) | Expansion | *Cryptotermes secundus* terminal branch |
| OG0002661 | Small ubiquitin-related modifier 3 (sumo3) | Expansion | *Coptotermes formosanus* terminal branch |
| OG0003454 | Corepressor interacting with RBPJ 1 (CIR1) | Expansion | *Zootermopsis nevadensis* terminal branch |
| OG0004273 | FAD-linked sulfhydryl oxidase ALR (Gfer) | Expansion | *Coptotermes formosanus* terminal branch |
| OG0004550 | Zinc finger protein 830 (znf830) | Expansion | *Cryptotermes secundus* terminal branch |

**Table S7.** List of single-copy orthologues that were under significant positive selection (*p* < 0.05) on ≥2 termite branches. The four termite branches were assigned as ‘foreground’ banches in this absREL analysis. Orthogroups that were not annotated were excluded.

| **Orthogroup** | **Putative annotation** | **Significant branches** |
| --- | --- | --- |
| OG0008616 | Glycogenin-1 (GYG1) | - *Cryptotermes secundus* terminal branch  - *Coptotermes formosanus* terminal branch  - *Coptotermes formosanus*/*Cryptotermes secundus* ancestral branch  - Termite stem branch |
| OG0006877 | Transmembrane protein 164 (Tmem164) | *- Zootermopsis nevadensis* terminal branch  - *Coptotermes formosanus* terminal branch  - Termite stem branch |
| OG0006995 | tRNA modification GTPase GTPBP3, mitochondrial (gtpbp3) | *- Zootermopsis nevadensis* terminal branch  - *Coptotermes formosanus* terminal branch  - Termite stem branch |
| OG0007009 | General transcription factor IIF subunit 2 (TfIIFbeta) | *- Zootermopsis nevadensis* terminal branch  - *Coptotermes formosanus* terminal branch  - Termite stem branch |
| OG0007542 | Phosphoacetylglucosamine mutase (PGM3) | *- Zootermopsis nevadensis* terminal branch  - *Cryptotermes secundus* terminal branch  - *Coptotermes formosanus* terminal branch |
| OG0007842 | Phospholipase A2 group XV (PLA2G15) | - *Cryptotermes secundus* terminal branch  - *Coptotermes formosanus* terminal branch  - Termite stem branch |
| OG0007964 | Protein angel homolog 2 (ANGEL2) | *- Zootermopsis nevadensis* terminal branch  - *Cryptotermes secundus* terminal branch  - *Coptotermes formosanus* terminal branch |
| OG0008181 | 39S ribosomal protein L40, mitochondrial (Mrpl40) | *- Zootermopsis nevadensis* terminal branch  - *Cryptotermes secundus* terminal branch  - Termite stem branch |
| OG0008245 | Ufm1-specific protease 1 (Ufsp1) | *- Zootermopsis nevadensis* terminal branch  - *Cryptotermes secundus* terminal branch  - Termite stem branch |
| OG0008527 | Transmembrane protein 184B (TMEM184B) | *- Zootermopsis nevadensis* terminal branch  - *Cryptotermes secundus* terminal branch  - *Coptotermes formosanus*/*Cryptotermes secundus* ancestral branch |
| OG0008710 | Ras-related GTP-binding protein C (Rragc) | *- Zootermopsis nevadensis* terminal branch  - *Cryptotermes secundus* terminal branch  - *Coptotermes formosanus* terminal branch |
| OG0006807 | Peptidyl-prolyl cis-trans isomerase (FKBP2) | - *Cryptotermes secundus* terminal branch  - Termite stem branch |
| OG0006897 | Succinate-semialdehyde dehydrogenase, mitochondrial (ALDH5A1) | *- Zootermopsis nevadensis* terminal branch  - Termite stem branch |
| OG0006947 | Short coiled-coil protein homolog (Bm1_04115) | - *Cryptotermes secundus* terminal branch  - *Coptotermes formosanus* terminal branch |
| OG0007013 | Vesicle transport through interaction with t-SNAREs homolog 1A (VTI1A) | *- Zootermopsis nevadensis* terminal branch  - *Coptotermes formosanus* terminal branch |
| OG0007021 | Beta-taxilin (TXLNB) | - *Coptotermes formosanus*/*Cryptotermes secundus* ancestral branch  - Termite stem branch |
| OG0007071 | Arginine/serine-rich coiled-coil protein 2 (rsrc2) | *- Zootermopsis nevadensis* terminal branch  - *Cryptotermes secundus* terminal branch |
| OG0007118 | Larval cuticle protein A1A | - *Coptotermes formosanus* terminal branch  - *Coptotermes formosanus*/*Cryptotermes secundus* ancestral branch |
| OG0007195 | Eukaryotic peptide chain release factor subunit 1 (eRF1) | - *Cryptotermes secundus* terminal branch  - Termite stem branch |
| OG0007277 | Vacuole membrane protein 1 (vmp1) | *- Zootermopsis nevadensis* terminal branch  - *Cryptotermes secundus* terminal branch |
| OG0007362 | Tryptophan 2,3-dioxygenase | *- Zootermopsis nevadensis* terminal branch  - *Cryptotermes secundus* terminal branch |
| OG0007424 | Probable 28S ribosomal protein S26, mitochondrial (mRpS26) | - *Coptotermes formosanus* terminal branch  - Termite stem branch |
| OG0007439 | Vacuolar-sorting protein (SNF8) | - *Cryptotermes secundus* terminal branch  - Termite stem branch |
| OG0007444 | Adapter molecule (Crk) | *- Zootermopsis nevadensis* terminal branch  - *Coptotermes formosanus* terminal branch |
| OG0007498 | 60S acidic ribosomal protein P0 (RpLP0) | - *Coptotermes formosanus* terminal branch  - *Coptotermes formosanus*/*Cryptotermes secundus* ancestral branch |
| OG0007572 | 28S ribosomal protein S14, mitochondrial (MRPS14) | *- Zootermopsis nevadensis* terminal branch  - *Coptotermes formosanus* terminal branch |
| OG0007648 | Protein DEK (Dek) | - *Cryptotermes secundus* terminal branch  - Termite stem branch |
| OG0007669 | Partner of Y14 and mago (Pym) | *- Zootermopsis nevadensis* terminal branch  - *Coptotermes formosanus* terminal branch |
| OG0007688 | Probable cardiolipin synthase (CMP-forming) (CLS) | *- Zootermopsis nevadensis* terminal branch  - *Cryptotermes secundus* terminal branch |
| OG0007757 | Insulin-like growth factor 2 mRNA-binding protein 1 (IGF2BP1) | - *Cryptotermes secundus* terminal branch  - Termite stem branch |
| OG0007814 | Endoplasmic reticulum resident protein 29 (Erp29) | - *Coptotermes formosanus* terminal branch  - *Coptotermes formosanus*/*Cryptotermes secundus* ancestral branch |
| OG0007858 | Protein max (MAX) | - *Coptotermes formosanus* terminal branch  - Termite stem branch |
| OG0007922 | Ankyrin repeat and LEM domain-containing protein 2 (Ankle2) | *- Zootermopsis nevadensis* terminal branch  - *Coptotermes formosanus*/*Cryptotermes secundus* ancestral branch |
| OG0007965 | Nurim homolog (nrm) | - *Cryptotermes secundus* terminal branch  - *Coptotermes formosanus* terminal branch |
| OG0007990 | Polymerase delta-interacting protein 2 (Poldip2) | *- Zootermopsis nevadensis* terminal branch  - *Coptotermes formosanus*/*Cryptotermes secundus* ancestral branch |
| OG0008005 | Transcriptional repressor scratch 1 (Scrt1) | - *Cryptotermes secundus* terminal branch  - Termite stem branch |
| OG0008006 | Protein Skeletor, isoforms B/C (Skeletor) | *- Zootermopsis nevadensis* terminal branch  - *Coptotermes formosanus* terminal branch |
| OG0008027 | SET domain-containing protein SmydA-8, isoform B (SmydA-8) | - *Coptotermes formosanus* terminal branch  - *Coptotermes formosanus*/*Cryptotermes secundus* ancestral branch |
| OG0008055 | JNK1/MAPK8-associated membrane protein (JKAMP) | *- Zootermopsis nevadensis* terminal branch  - *Cryptotermes secundus* terminal branch |
| OG0008106 | Proteasome subunit beta type-7 (Psmb7) | *- Zootermopsis nevadensis* terminal branch  - *Cryptotermes secundus* terminal branch |
| OG0008133 | Cuticle protein 19 | *- Zootermopsis nevadensis* terminal branch  - *Cryptotermes secundus* terminal branch |
| OG0008182 | Vesicle-associated membrane protein 7 (Vamp7) | - *Cryptotermes secundus* terminal branch  - *Coptotermes formosanus* terminal branch |
| OG0008193 | Acidic fibroblast growth factor intracellular-binding protein (Fibp) | *- Zootermopsis nevadensis* terminal branch  - *Coptotermes formosanus* terminal branch |
| OG0008214 | CKLF-like MARVEL transmembrane domain-containing protein 4 (Cmtm4) | *- Zootermopsis nevadensis* terminal branch  - *Cryptotermes secundus* terminal branch |
| OG0008358 | V-type proton ATPase subunit d | *- Zootermopsis nevadensis* terminal branch  - *Coptotermes formosanus* terminal branch |
| OG0008390 | RING finger protein 121 (rnf121) | - *Coptotermes formosanus* terminal branch  - Termite stem branch |
| OG0008434 | Hydroxymethylglutaryl-CoA lyase, mitochondrial (HMGCL) | - *Coptotermes formosanus* terminal branch  - Termite stem branch |
| OG0008575 | CDP-diacylglycerol--inositol 3-phosphatidyltransferase (Pis) | - *Cryptotermes secundus* terminal branch  - Termite stem branch |
| OG0008609 | Cyclic AMP-dependent transcription factor ATF-6 alpha (Atf6) | *- Zootermopsis nevadensis* terminal branch  - *Coptotermes formosanus*/*Cryptotermes secundus* ancestral branch |
| OG0008624 | NADH dehydrogenase [ubiquinone] flavoprotein 2, mitochondrial (NDUFV2) | *- Zootermopsis nevadensis* terminal branch  - *Cryptotermes secundus* terminal branch |
| OG0008629 | Nucleic acid dioxygenase (Alkbh1) | - *Coptotermes formosanus* terminal branch  - Termite stem branch |
| OG0008653 | TM2 domain-containing protein almondex (amx) | - *Coptotermes formosanus* terminal branch  - Stem termite branch |
| OG0008662 | CDK5RAP1-like protein (CG6345) | - *Cryptotermes secundus* terminal branch  - *Coptotermes formosanus* terminal branch |
| OG0008669 | Beta-1,4-glucuronyltransferase 1 (b4gat1) | *- Zootermopsis nevadensis* terminal branch  - *Coptotermes formosanus* terminal branch |
| OG0008715 | Ribonuclease P protein subunit p25-like protein (RPP25L) | *- Zootermopsis nevadensis* terminal branch  - Termite stem branch |

**Supplemental Figures**

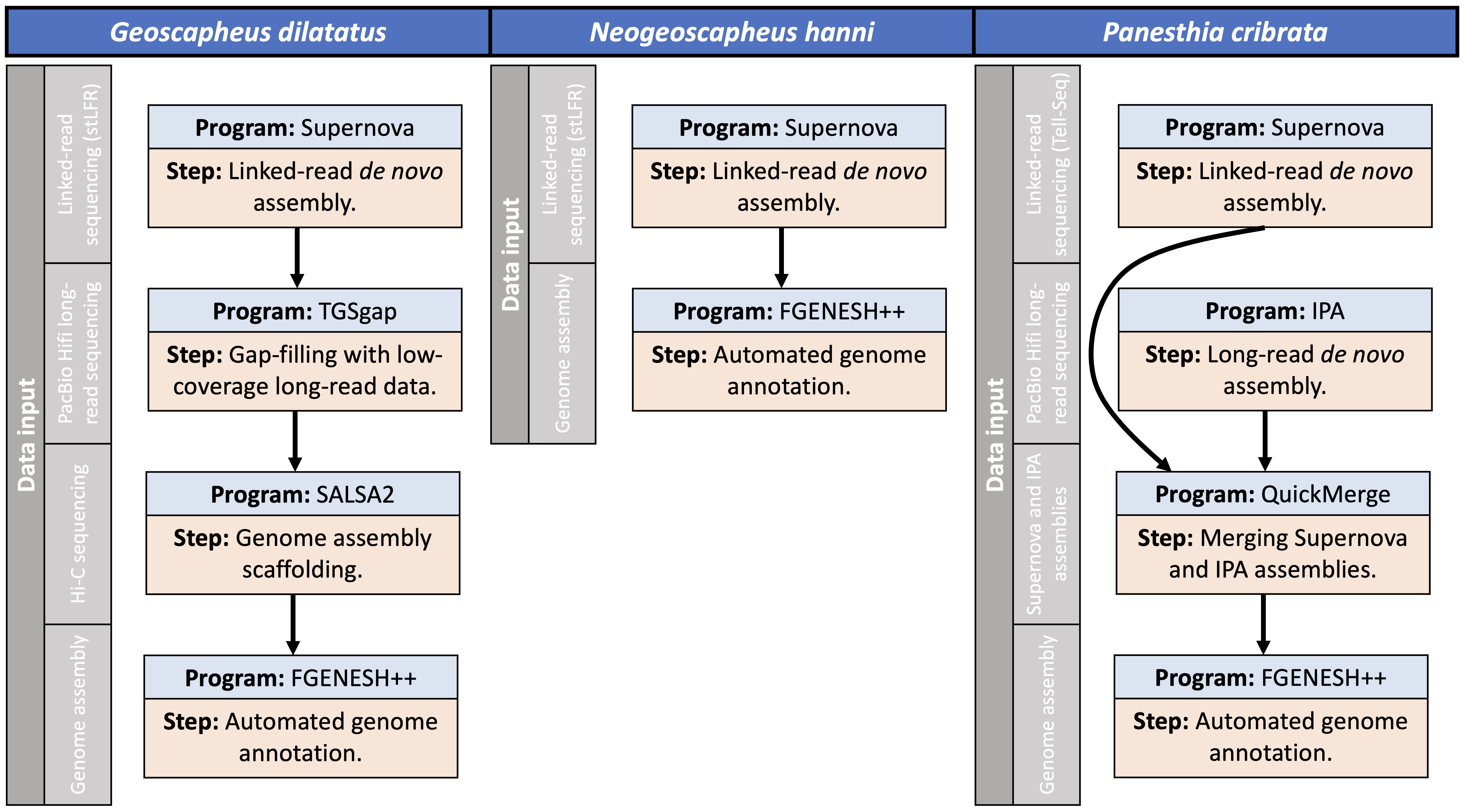

**Figure S1.** Genome assembly/annotation pipelines for the three cockroach species.

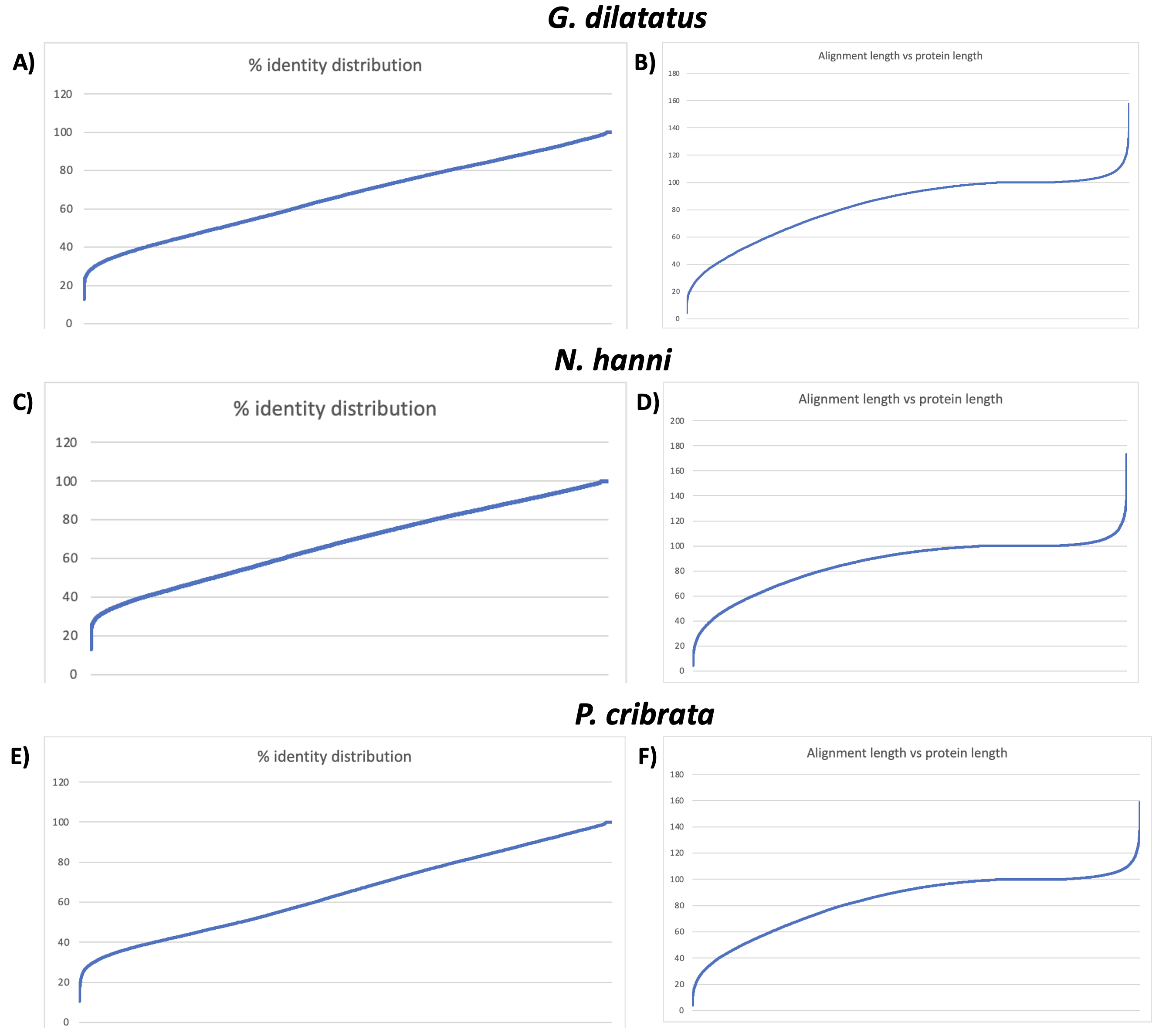

**Figure S2.** Sequence identity distribution when BLASTing the annotated genes against the insect protein database for *Geoscapheus dilatatus* (A), *Neogeoscapheus hanni* (C) and *Panesthia cribrata* (E); and the alignment length of the top BLAST hit for each gene for *G. dilatatus* (B), *N. hanni* (D) and *P. cribrata* (F). The average resultant sequence identity was 66.4%, 67.9%, and 64.7% for *G. dilatatus*, *N. hanni*, and *P. cribrata*, respectively, and the proportion of BLAST hits whereby the alignment length was >50% of the annotated protein length was 88.27%, 91.9%, and 88.9% for *G. dilatatus*, *N. hanni*, and *P. cribrata*, respectively.

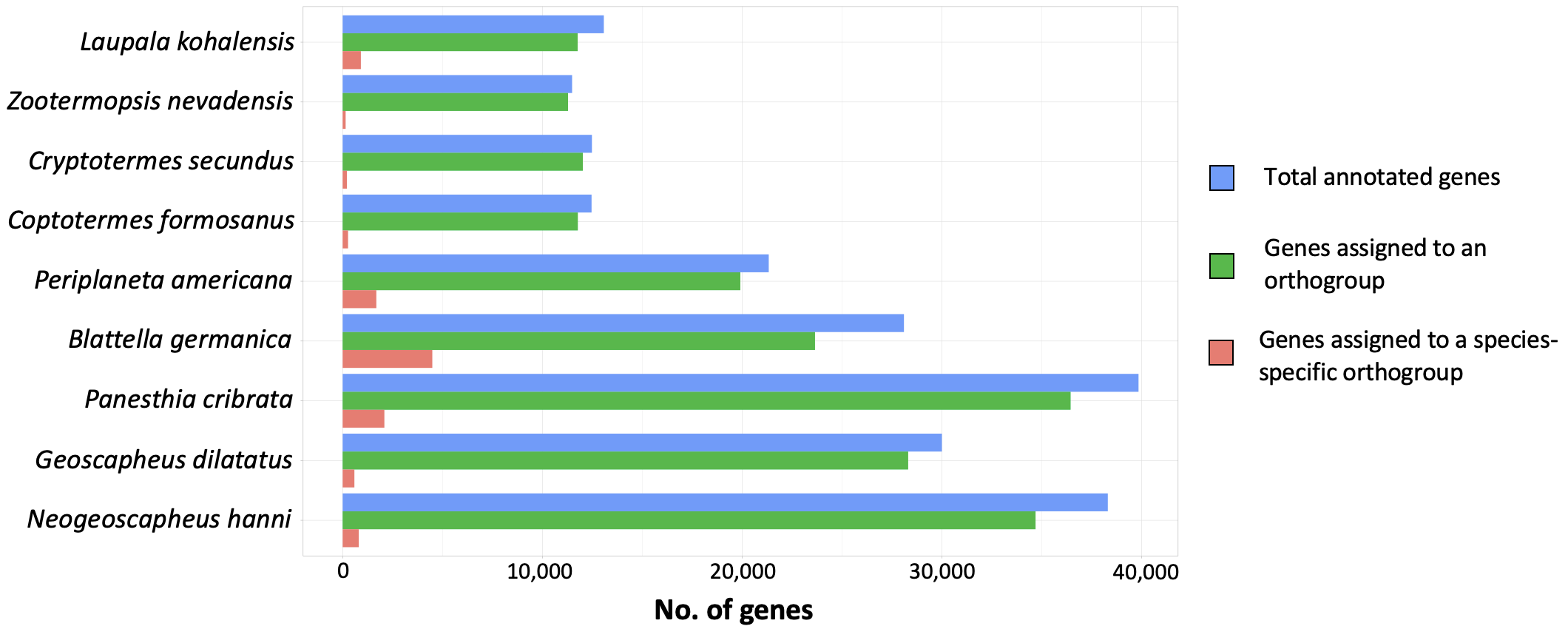

**Figure S3.** Orthogroup clustering results used in the subsequent comparative genomic analyses.

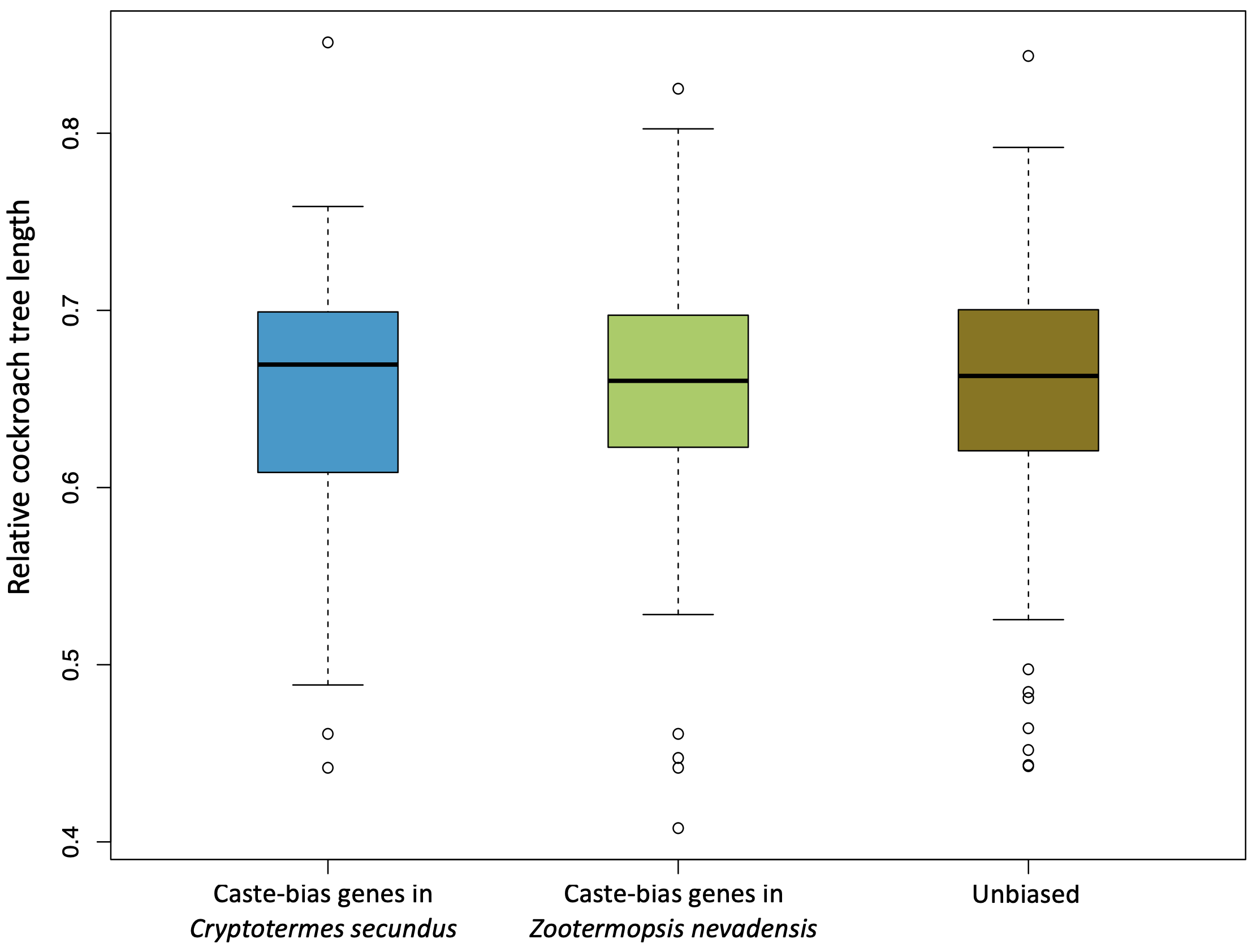

**Figure S4.** Distributions of relative cockroach tree lengths (cockroach tree length divided by the whole tree length) for unbiased and caste-biased genes identified in *Cryptotermes secundus* and *Zootermopsis nevadensis*. There were no significant differences between relative cockroach tree lengths across the biased and unbiased genes based on Mann-Whitney U tests.

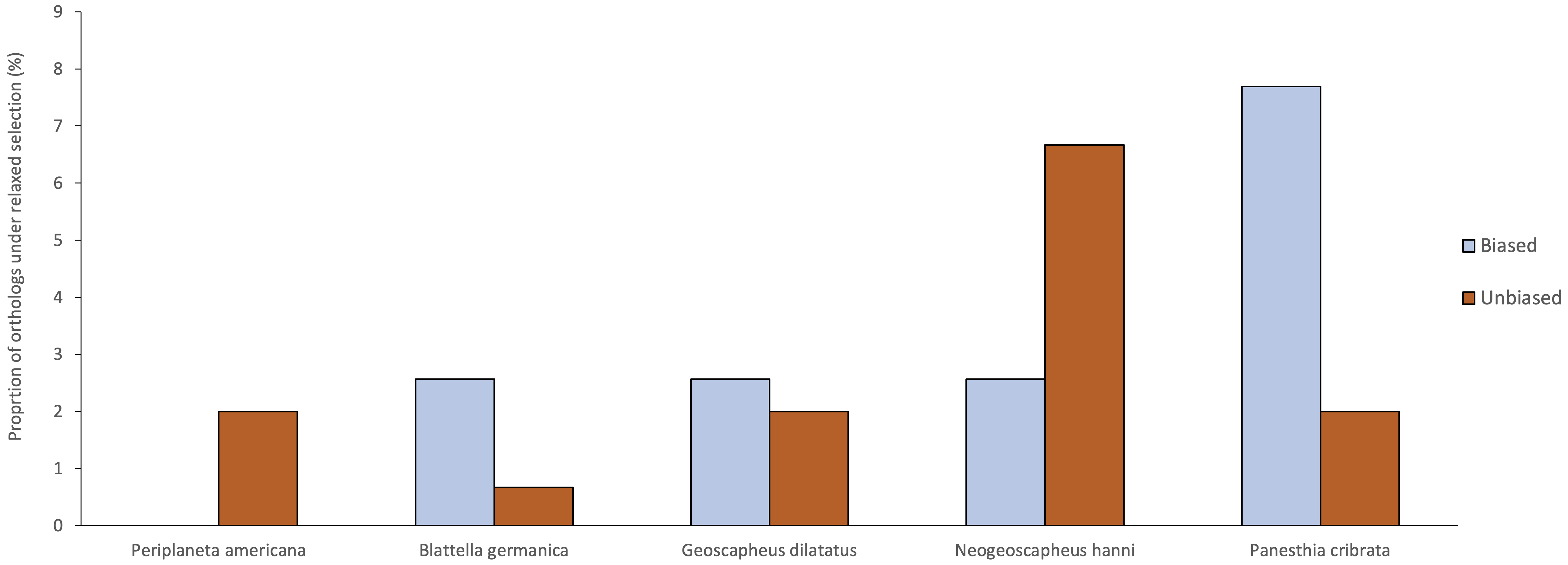

**Figure S5.** Proportion of caste-biased (blue) and unbiased (orange) genes from Harrison et al. (2021) and Maekawa et al. (2022) identified in our data set under relaxed selection in the five cockroach terminal branches. The biased (*n* = 39) and unbiased (*n* = 150) genes were only considered if they were biased/unbiased in both *Zootermopsis nevadensis* and *Cryptotermes secundus*. There were no significant differences between the unbiased and biased genes under relaxed selection for any of the five species based on chi-square tests. This analysis was included to investigate the hypothesis that caste-biased genes were co-opted from genes that were already under relaxed selection.

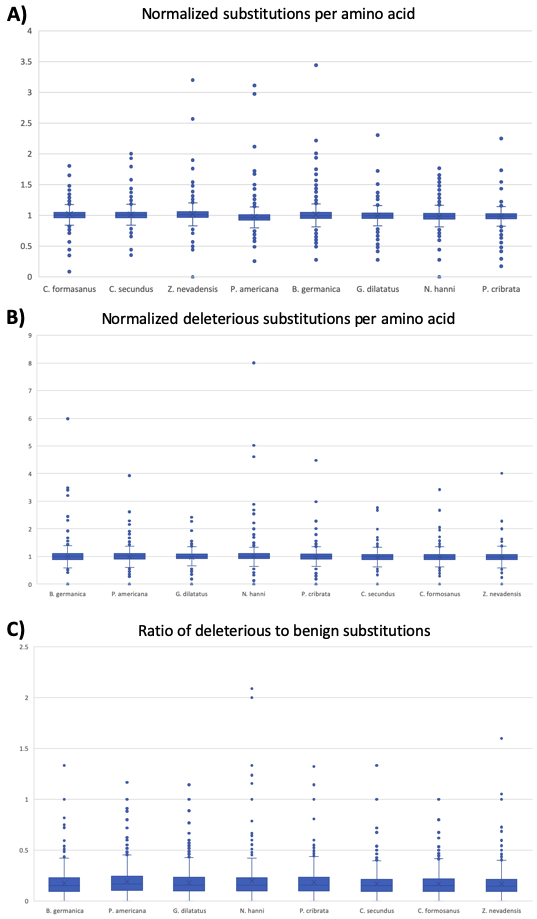

**Figure S6.** A) Number of substitutions per amino acid for each species when compared with the outgroup, *L. kohalensis* (values have been normalized for visualization). B) Number of mildly and moderately deleterious substitutions (as determined using PROVEAN) per amino acid for each species when compared with the outgroup (values have been normalized for visualization). C) Ratio of deleterious to benign amino acid substitutions for each species.

**Supplemental Methods**

*DNA extractions:*

Collected whole specimens of *P. cribrata*, *G. dilatatus*, and *N. hanni* were stored at -20°C for 5 min, after which one middle leg was removed; individuals were then returned to the freezer. Muscle tissue was removed from the leg of one individual from each species and crushed using a sterile pestle in a 1.5 mL Eppendorf tube containing 600 µL of SDS Buffer and 20 µL of Proteinase K (Roche, Switzerland), then incubated at 56°C overnight. Following tissue lysis, 5 µL of RNAse A (20 mg/mL) was added, followed by incubation at 37°C for 1 h. We extracted DNA using phenol/chloroform/isoamyl alcohol (600 µL), with rotation for 15 min, followed by centrifugation at 16,000 g. The aqueous phase was taken and placed into a new tube, to which 600 µL of chloroform was added, followed by rotation and centrifugation as described above. The aqueous phase was then placed in a new tube with 40 µL of 3 M NaAc buffer (pH 5.5) and 620 µL of ice-cold isopropanol, and centrifuged. The pellet was washed with 70% EtOH. DNA concentration was measured using Qubit fluorometric quantitation (ThermoFisher, USA), and DNA quality was visualized using 1% agarose gel electrophoresis.

*Library preparation:*

TELL-seq data were generated by the Australian Genome Research Facility. TELL-seq libraries were constructed using a TELL-seq WGS Library Prep Kit (Universal Sequencing Technology). DNA extracted from *P. cribrata* was incubated with ~8 million TELL beads for barcoding in a 0.2 mL PCR tube according to the manufacturer’s protocol (performed by AGRF). Each TELL bead comprises 50,000 copies of at least one barcode sequence conjugated to its surface. Following barcoding, an eight-cycle amplification step was carried out in order to generate libraries for sequencing. stLFR libraries for *G. dilatatus* and *N. hanni* were prepared by BGI Shenzen, using the procedure outlined by Wang et al. (2019).

Hi-C library preparation was performed using the Arima Hi-C Plus kit (Arima, USA) following the manufacturer’s protocol for large animal tissue, using the *DpnII* and *HinfI* enzymes. Up to 50 mg of fresh cockroach muscle tissue was flash frozen and pulverized using liquid nitrogen. A crosslinking buffer containing 1% formaldehyde was added to the pulverized tissue to crosslink DNA. We used 500 ng of DNA for Hi-C library preparation. The final library preparation for sequencing was performed using the KAPA Hyper Prep kit following the protocol detailed in the Arima Hi-C Plus kit. The libraries were quantified using a Qubit fluorometer (ThermoFisher, USA) and Bioanalyzer (Agilent, USA). In order to estimate the number of available Hi-C reads in the library, the library was first sequenced on the Illumina Miseq platform (Illumina, USA) using a Miseq V2 Nano flow cell with 2×150 bp specification. The sequencing data were then processed using qc3C (DeMaere and Darling, 2021). The qc3C software assessed the Hi-C library quality by calculating proximity ligation events which create k-mers that would not naturally occur in the sample. Based on the qc3C result, we estimated the sequencing data needed for the library. Sequencing was carried out on the Novaseq platform (Illumina, USA) using Novaseq S4 flow cell 2×150 bp at Novogene (USA).

The PacBio HiFi library preparation was performed using the SMRTbell® Express Template Prep Kit 2.0 following the manufacturer’s instructions (Pacific Biosciences). The library was sequenced on the PacBio Sequel II platform using one SMRT Sequel II cell at the Okinawa Institute of Science and Technology Graduate University (OIST). Subreads and circular consensus sequencing (CCS) reads were generated using the SMRTLink software with default settings. The Sequel II data set was filtered after CCS correction to retain only reads of HiFi quality standards (>Q20). We obtained a total of 13.3 Gigabases of HiFi reads.

**Supplemental Results**

*Analysis of genetic load and its caveats:*

As a conservative approach, we only divided derived mutations into putatively ‘benign’ and ‘deleterious’. We acknowledge that this coarse assignment of mutations, and the lack of dominance information, does not fully capture the complexities of genetic load effects and purging across mutations of varying degrees of deleteriousness (Dussex et al., 2023). Further, in our analysis, we consider deleterious mutations relative to the cricket genome, which does not represent a mutant-free reference genotype; this is a ubiquitous limitation in investigations of genetic load (Agrawal & Whitlock, 2012).

We acknowledge issues of non-independence in our phylogenetic approach to investigate genetic load; several ancestral branches are considered multiple times in the species comparisons. For several reasons, however, we believe that these issues have a negligible effect on our inferences. First, a large proportion of the genes under relaxed selection occur at the terminal branches (Figure 1c), hence the effects of *N*_e_ and the potential accumulation of deleterious alleles in ancestral branches appears to be a less prominent issue. The MRCA for the three termite species analysed is ~107 million years ago (Evangelista et al., 2019), which is ample time for genetic load to accumulate after a dramatic shift in *N*_e_. Second, although we cannot rule out that the genetic load and relaxed selection patterns identified in the three termite species is a clade effect due to their common ancestry (i.e., representing one data point instead of three), we note that *Periplaneta americana*, the sister species to termites among the taxa considered, has a much lower proportion of orthologues under relaxed selection than the termites, yet harbours a significantly higher or comparable number of putative deleterious alleles (Table S5).

*Positive selection:*

For the aBSREL positive selection analyses focusing on the four termite branches (i.e., assigned as the ‘foreground’ branches), the best-fitting models accounting for multinucleotide mutations varied across the different orthologues based on AICc scores; the standard model was best-fitting for 43.6% of orthologues, the multiple-hits ‘Double’ model was best-fitting for 50.6% of orthologues, and the multiple hits ‘Double+Triple’ model was best-fitting for 5.8% of orthologues. We retained analyses implementing the best-fitting model for each orthologue. For the aBSREL positive selection analyses considering the larger subset of branches (i.e., assigned as ‘foreground’ branches), including the cockroach terminal branches, the standard model was best-fitting for 43.5% of orthologues, the multiple-hits ‘Double’ model was best-fitting for 50.2% of orthologues, and the multiple hits ‘Double+Triple’ model was best-fitting for 6.3% of orthologues.

In our analyses that allowed a separate *d_N_/d_S_* on terminal branches leading to cockroach and termite taxa, we detected 28.24% (*n* = 421) of single-copy orthologues as being under positive selection on at least one branch (after correcting for multiple testing). The number of orthologues under positive selection varied between 1.1% and 9.1% across the terminal branches (Figure 2a), with the clade comprising *P. cribrata*, *G. dilatatus*, and *N. hanni* displaying the highest number. When focusing on specific genes that might be under positive selection in the termites, by allowing a separate *d_N_/d_S_* on termite lineages, we found evidence for 3.76% (*n* = 56) of the 1491 single-copy orthologues being under positive selection on at least two of the termite branches (Table S6). We inferred 4.6%, 7.6%, and 4.6% of the orthologues to be under positive selection on the terminal branches leading to *Zootermopsis nevadensis*, *Coptotermes formosanus*, and *Cryptotermes secundus*, respectively.

*Gene expansions and contractions:*

Across all branches in the tree comprising the eight cockroach and termite species, 8538 orthogroups displayed size changes (Figure 2b), 1134 orthogroups of which exhibited significant contractions or expansions. Of these, 40 displayed significant (*p* < 4×10-5) expansions (*n* = 35) or contractions (*n* = 5) on the termite branches (Table S7), all of which were restricted to a single species (i.e., 8 in *Zootermopsis nevadensis*, 15 in *Coptotermes formosanus*, and 17 in *Cryptotermes secundus*).

**Supplemental References**

Agrawal, A. F., & Whitlock, M. C. (2012). Mutation load: the fitness of individuals in populations where deleterious alleles are abundant. *Annual Review of Ecology, Evolution, and Systematics*, *43*, 115-135.

DeMaere, M. Z., & Darling, A. E. (2021). qc3C: reference-free quality control for Hi-C sequencing data. *PLoS Computational Biology*, *17*, e1008839.

Dussex, N., Morales, H. E., Grossen, C., Dalén, L., & van Oosterhout, C. (2023). Purging and accumulation of genetic load in conservation. *Trends in Ecology & Evolution*, *10*, 961-969.

Evangelista, D. A., Wipfler, B., Béthoux, O., Donath, A., Fujita, M., Kohli, M. K., ... & Simon, S. (2019). An integrative phylogenomic approach illuminates the evolutionary history of cockroaches and termites (Blattodea). *Proceedings of the Royal Society B*, *286*, 20182076.

Harrison, M. C., Chernyshova, A. M., & Thompson, G. J. (2021). No obvious transcriptome‐wide signature of indirect selection in termites. *Journal of Evolutionary Biology*, *34*, 403-415.

Maekawa, K., Hayashi, Y., & Lo, N. (2022). Termite sociogenomics: evolution and regulation of caste-specific expressed genes. *Current Opinion in Insect Science*, *50*, 100880.

Wang, O., Chin, R., Cheng, X., Wu, M. K. Y., Mao, Q., Tang, J., ... & Peters, B. A. (2019). Efficient and unique cobarcoding of second-generation sequencing reads from long DNA molecules enabling cost-effective and accurate sequencing, haplotyping, and de novo assembly. *Genome Research*, *29*, 798-808.
